# Ligand-Mediated Endocytosis Is Regulated in A Sexually Dimorphic Way in Osteocytes *in vivo*

**DOI:** 10.1101/2024.08.18.608500

**Authors:** Melia Matthews, Alexander Saffari, Nuzhat Mukul, Lanlan Hai, Nada Naguib, Ulirich B. Wiesner, Karl J. Lewis

**Affiliations:** Department of Biomedical Engineering, Cornell University, 237 Tower Rd, Ithaca NY 14850, New York, USA; Department of Materials Science and Engineering, Cornell University, Bard Hall 210, Ithaca NY 14850, USA; Kavli Institute at Cornell for Nanoscale Science, Cornell University, Ithaca Y 14853, USA; Department of Design Tech, Cornell University, Ithaca NY 14853, USA

**Keywords:** osteocyte, fluorescent nanoparticles, integrin, endocytosis

## Abstract

Endocytosis is a critical cellular process involved in many physiological functions. Most research on endocytosis has been performed *in vitro*, however, understanding this process *in vivo* is necessary in tissues like bone that have a unique 3D extra-cellular matrix. Here, we present a live-cell study of endocytosis in osteocytes, mechanosensory cells embedded in mouse bone. We visualized real-time fluorescent nanoparticle uptake and trafficking in osteocytes by using intravital imaging combined with multiphoton microscopy within living animals. We applied pharmacologic inhibitors to distinguish between general and receptor-specific endocytosis pathways *in vivo*. Our findings reveal rapid nanoparticle uptake in osteocytes, with marked differences in the timescale and pattern of uptake depending on nanoparticle surface functionality. We also discovered differences in dynamin-dependent endocytosis in osteocytes between male and female animals. These results offer the first *in vivo* derived insights into how osteocytes take up materials and provide new evidence for chemically altering receptor-mediated endocytosis in live bone tissue.

## 1. Introduction

Endocytosis is a fundamental cellular process that enables cells to sense and respond to their external environment by internalizing molecules from the extracellular matrix. It is crucial for maintaining the function of membrane-bound proteins (1), receptor metabolism (2, 3), regulating neurotransmitter uptake (4–6) and regulation of mechanotransduction machinery (7, 8). Through endocytosis, cells modulate the surface availability of transmembrane proteins, facilitating precise spatiotemporal control of signaling pathways (9–12). This process underpins essential cellular functions, including migration and mechanotransduction (1, 2). Despite the central role of endocytosis in maintaining cellular homeostasis, many aspects of the molecular mechanisms governing different endocytic pathways and their physiological implications remain poorly understood.

In mammalian cells, endocytosis occurs via multiple pathways (8, 13–17). These pathways have been extensively characterized *in vitro*, often using pharmacological inhibitors that disrupt specific stages of vesicle formation or receptor internalization. For example, clathrin-mediated endocytosis is critical for receptor uptake in several cell types, including fibroblasts and neurons (18, 19). Cholesterol-dependent endocytic pathways, such as those involved in lipid raft-mediated uptake, have also been identified (20, 21). Clathrin-mediated pathways can be inhibited by dynamin blockers (19) while cholesterol-dependent pathways can be disrupted by agents that deplete membrane cholesterol (22, 23). *In vitro* studies have elucidated much of the machinery and dynamics of endocytosis, but they are limited by the inability to fully replicate the complexities of living systems, where cell-cell interactions, mechanical forces, and tissue architecture play crucial regulatory roles (24, 25). Additionally, culture conditions can artificially influence membrane tension and alter endocytic activity, biasing specific pathways (26, 27). Thus there is a critical need for *in vivo* experimentation to more fully understand the implications of endocytic pathways in cell homeostasis.

The development of advanced imaging modalities, particularly multiphoton microscopy, has provided a pathway to overcome some experimental limitations by enabling high-resolution, real-time visualization of endocytosis *in vivo* (25). Consequently, researchers can explore endocytosis in the context of intact tissue environments. The scope of these *in vivo* studies remains narrow, however. Most *in vivo* endocytosis exploration has focused primarily on kidney and salivary gland model organ systems using fluorescently labeled dextran as a marker of bulk endocytosis (28–30). The limited scope of *in vivo* endocytosis research is in large part due to the technical challenges associated with new model development. Notably, there has been little to no investigation of endocytosis in bone tissue, a dynamic and mechanosensitive environment where there is relatively limited fundamental understanding of surface protein dynamics and metabolism.

Bone is a highly mechanosensitive tissue. Wolff’s Law, originally reported in the 19th century, states that bone tissue changes according to the mechanical loads it experiences (31). Indeed, bone relies on force for healthy form and function (32–34). Tissue adaptation is facilitated by bone specific cells; bone building osteoblasts and bone resorbing osteoclasts. Recent studies suggest a significant role for endocytosis in osteoblast activity; for example, the conditional deletion of a caveolin-dependent endocytosis component in osteoblasts enhances bone formation and improves tissue mechanical properties (35, 36). Critically, these studies have been performed almost universally *in vitro*. Moreover, membrane trafficking pathways have not been interrogated in an integral third cell type within bone, the osteocyte.

Osteocytes sense and translate mechanical stimulation into biochemical signals to control the actions of osteoblasts and osteoclasts (37, 38). These dendritic cells are embedded in mineralized bone matrix and serve as the resident mechanosensor within the tissue (39). Osteocyte function is critical to bone homeostasis. However, many features of their basic function have not been studied, including transmembrane protein endocytosis. The dearth of information about osteocyte biology exists because they are very challenging cells to study. Primary osteocytes are difficult to remove from bone tissue and culture *in vitro* (40). Immortalized cell lines recapitulate gene expression and differentiation of osteocytes, but do not reliably mirror the unique 3D mineralized environment within bone (41). Studying osteocyte activity *in vivo* is also difficult due to the technical challenge of visualizing cells embedded within the highly optically scattering bone matrix. Classical imaging methodologies do not allow enough depth penetration or resolution to interrogate the dynamic activity of osteocytes in their native environment. Novel methodologies and techniques are therefore needed to answer cellular level questions in living bone, including the role of endocytosis *in vivo*.

Our group has developed of an intravital imaging platform that allows the investigation of osteocyte biology *in vivo*, maintaining the intact vasculature and microenvironment of bone tissue (42). This platform leverages ultrasmall (*<* 7nm), highly fluorescent nanoparticles termed C’Dots (42, 43). These nanoparticles are synthesized in water and have a silica core with a poly(ethylene glycol) (PEG) shell (43, 44). C’Dots exhibit rapid cellular internalization and transient retention, making them ideal for tracking endocytosis dynamics in real time. C’Dots can also be surface functionalized to target specific cellular uptake pathways, providing a unique opportunity to dissect endocytic mechanisms *in vivo*. Furthermore, this system allows for localized pharmacological manipulation, enabling precise perturbation of endocytic pathways within the bone microenvironment.

In this study, we utilized three different types of C’Dot surface functionalities to investigate the mechanisms of nanoparticle uptake and clearance in osteocytes *in vivo*. They include untargeted (i.e., PEGylated), integrin-targeted, and membrane-penetrating C’Dots. We hypothesized that osteocyte endocytosis governs the internalization of these nanoparticles. We explore this hypothesis by applying pharmacological inhibitors of clathrin-mediated endocytosis and macropinocytosis (i.e., Dyngo4a and Methyl-β-cyclodextrin, respectively) to disrupt these pathways. By examining the effects of these inhibitors on osteocyte nanoparticle uptake and clearance, we aim to establish a foundational understanding of endocytic dynamics in osteocytes within the intact bone environment and how it can be altered by pharmacological compounds. This work represents methodological advances in studying endocytosis in musculoskeletal tissue and provides critical insights into the cellular biology of bone.

## 2. Results

### 2.1 C’Dots are rapidly internalized by osteocytes *in vivo*

We developed a method to deliver fluorescent nanoparticles (C’Dots) to osteocytes *in vivo* (42). A 15 *µ*L subcutaneous injection of a 10 *µ*M C’Dot solution over the third metatarsal (MT3) of a mouse consistently provided clear images of the osteocyte microenvironment (**Figure 1A**). In **Figure 1B**), a representative image shows the red fluorescent Cy5 C’Dot signal present in all cells within the field of view. Some cells displayed uniform fluorescence throughout, while others exhibited distinct subcellular clusters of signal (**Figure 1C–F**). C’Dots were also rapidly cleared from the cells over time (**Figure 1H**). Notably, C’Dots were clustered into subcellular structures beyond the osteocyte cell bodies (**Figure 1G**). By combining a short C’Dot incubation period with optical zoom and high frame averaging, we were able to reproducibly visualize the dendritic network of osteocytes (**Figure 2A-C**). Together, these observations raise the question of whether C’Dots are internalized by osteocytes or simply adhere to the cell membrane.

**Fig. 1.**
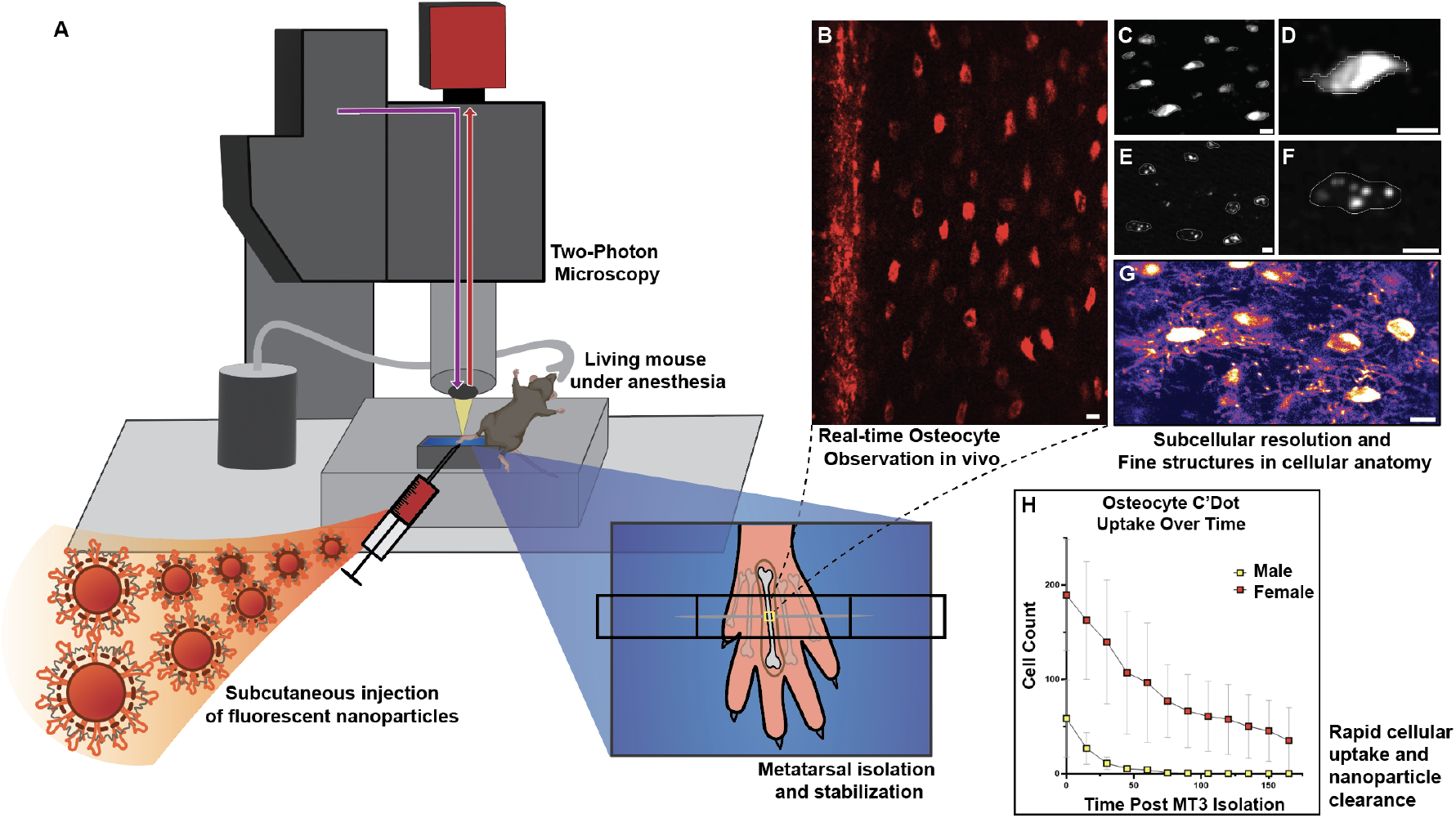
Graphical abstract of experimental design for imaging intravital C’Dot uptake and clearance in bone with 2-photon microscopy. Fluorescent nanoparticles were exposed to bone and osteocytes through a local subcutaneous injection above the third metatarsal (MT3) bone in a mouse hind paw (A). After C’Dot incubation, the MT3 was surgically isolated and secured for multiphoton imaging. Example images of C’Dots within osteocytes are shown in panels B-G, highlighting the high resolution of our imaging system which enables subcellular localization. Images were analyzed for clearance kinetics of nanoparticle signal (H) as well as subcellular distribution of signal. Scale bars = 10*µ*m.

**Fig. 2.**
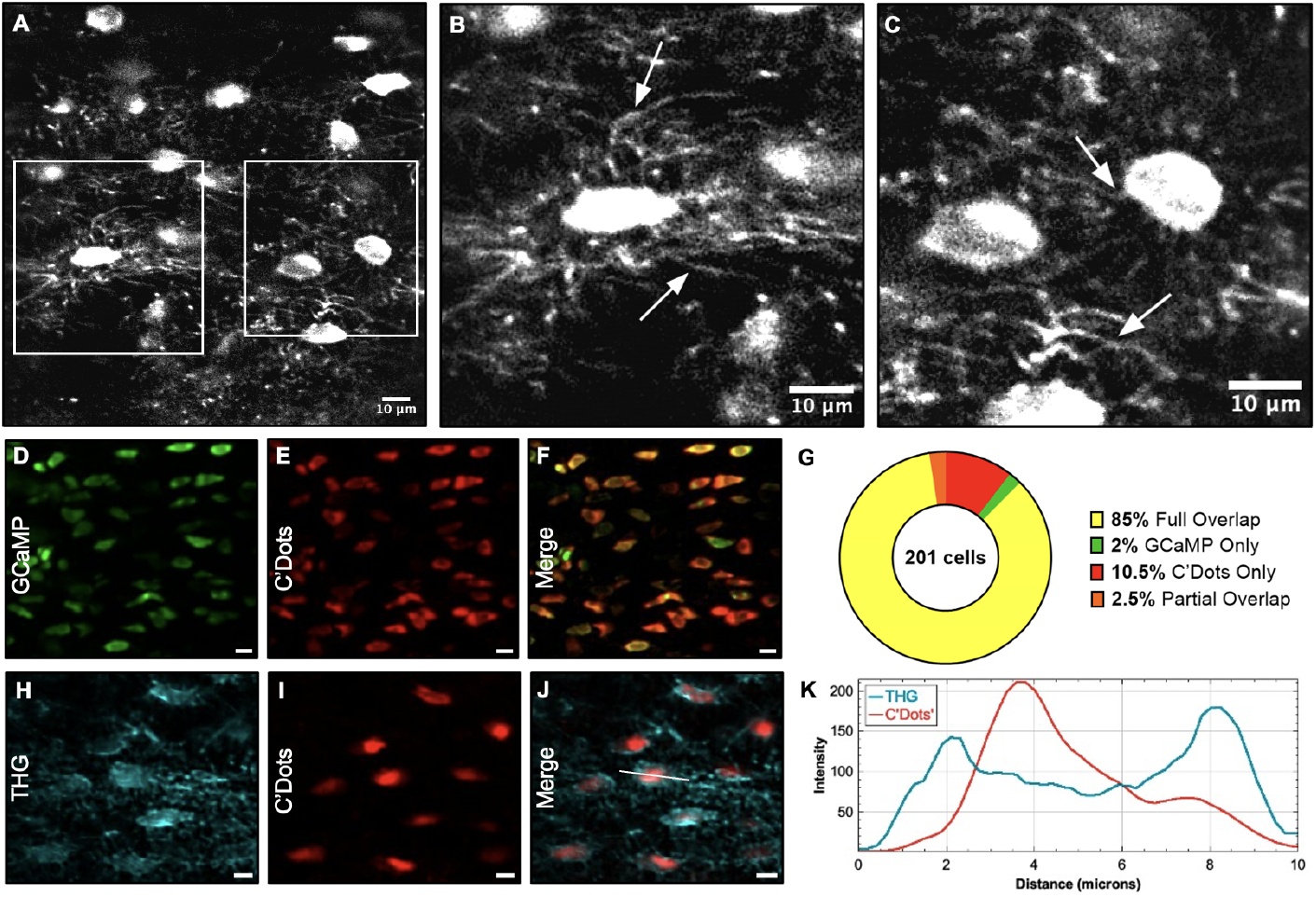
Osteocytes rapidly internalize subcutaneously injected C’Dots in vivo. A 2-minute incubation resulted in C’Dot uptake in vivo with dendrites clearly visible (A-C). To assess C’Dot and cell co-localization, genetically modified mice expressing GCaMP6f signal within osteocytes were locally injected with C’Dots for a 45-minute incubation before imaging. GCaMP signal (green) was collected using 920nm excitation and C’Dot signal (red) was collected using 1090nm excitation (D-F). Co-localization analysis on the overlap of signal showed a high percentage of cells with full or partial overlap with C’Dots (G). In another approach, wild-type mice were locally injected with C’Dots 45 minutes before imaging them with endogenous third harmonic generation (THG) signal (teal) using a 3-Photon microscope (H-J). Analysis of signal intensity across the linear profile of a representative cell (J, white line) is shown (K). The THG signal, which highlights material interfaces (i.e. matrix/fluid boundary), was external to C’Dot signal.

To that end, we introduced two co-localization approaches. First, we used a genetically modified mouse model expressing the calcium indicator GCaMP6f specifically in osteocytes (45). In these mice, GCaMP6f fluorescence is restricted to the intracellular cytosolic space (46). Subcutaneous injections of C’Dots showed a high degree of overlap between the red Cy5 C’Dot signal and the green GCaMP6f signal in osteocytes *in vivo* (**Figure 2D-F**). Untargeted PEG-C’Dots had complete signal overlap in 78% of cells, while integrin-targeted RGD-C’Dots showed similar overlap in 83% of cells (**Figure 2G, Supplemental Figure 1**). These results strongly suggest that C’Dots are primarily localized inside the osteocytes rather than on their external membrane.

In a second approach, we used three-photon microscopy with third harmonic generation (THG) imaging to further investigate C’Dot localization. THG, a dye-free signal generated at material interfaces like the cell membrane (47), was used to differentiate between intracellular and extracellular nanoparticle localization. Line intensity analysis across the diameter of each cell revealed that the Cy5 C’Dot signal was nested within the THG peaks, consistent with C’Dots localization predominantly inside the cell membrane (**Figure 2H-K, Supplemental Figure 1**). Together, these data provide strong evidence that C’Dots are internalized by osteocytes following subcutaneous injection, supporting the idea that osteocytes utilize endocytosis to take up nanoparticles.

### 2.2 C’Dot surface functionalization impacts uptake and clearance dynamics *in vivo*

To investigate the presence of distinct endocytic pathways in osteocytes, we examined the uptake of nanoparticles with different surface functionalization. We used three types of C’Dots to target specific uptake mechanisms in osteocytes: 1) Untargeted (i.e., purely PEGylated) PEG-C’Dots, serving as a control for non-receptormediated uptake; 2) integrin-targeting (cyclic RGD functionalized) RGD-C’Dots, to assess receptor-mediated uptake; and 3) TAT peptide functionalized TAT-C’Dots, which incorporate a cell-penetrating peptide derived from HIV, acting as a positive control for uptake independent of endocytosis (**Figure 3**). We designed experiments to observe both shortterm and extended nanoparticle dynamics within osteocytes. C’Dots were injected subcutaneously over the mouse MT3 and incubated for either 5 minutes (short incubation) or 45 minutes (long incubation) before imaging. We captured ten z-stacks over 30 minutes or 2.5 hours to assess nanoparticle kinetics for the short and long incubation times, respectively. At each time point, we quantified the total number of cells that had taken up C’Dot signal. In the short incubation study we used linear regressions to assess the initial cell uptake (y-intercept) and clearance (slope). In the long incubation study we compared the kinetics of C’Dot uptake and clearance between groups using 2-Way ANOVAs with multiple comparisons.

**Fig. 3.**
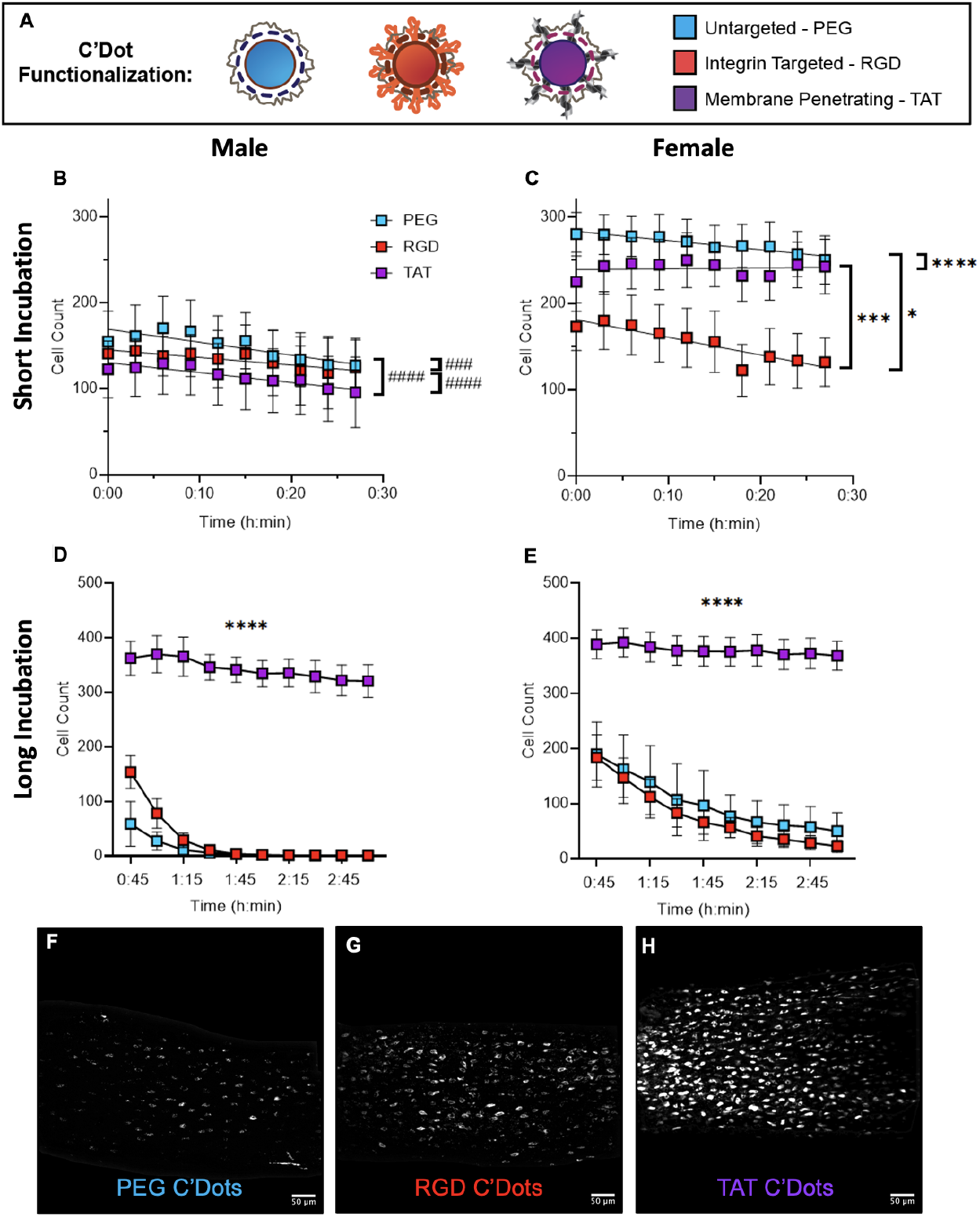
Functionalization of C’Dots with different cellular targeting peptides modulates their short and long term uptake and clearance. Three different C’Dots with distinct surface functionalization were tested (A). PEG-C’Dots are untargeted, RGD-C’Dots are integrin targeting with cyclic RGD motifs, and TAT-C’Dots are membrane penetrating, evading the endocytic pathway via functionalization with HIV viral peptides. Simple linear regression measures the impact of surface functionalization on short incubation (5 min.) based C’Dot uptake in osteocytes over time (B-C). Statistical measurements compare C’Dot surface functionalization to each other within each panel. A significant difference in slope is shown as *p≤0.05, **p≤0.01, ***p≤0.001, ****p≤0.0001. A significant difference in y-intercept is shown as #p≤0.05, ##p≤0.01, ###p≤0.001, ####p≤0.0001. Clearance curves after long incubation (45 min.) are also compared between surface functionalized particle groups with a 2-Way ANOVA (D-E). Representative z-projection images for each C’Dot type (as indicated) after a 45-minute incubation are shown in panels F-H. Error = SEM. Scale bars = 50*µ*m.

In the short incubation study, each C’Dot group exhibited distinct y-intercepts in male mice (**Figure 3A,B**). Overall, untargeted PEG-C’Dots showed slightly higher uptake compared to both RGD and TAT-C’Dots, suggesting faster internalization of non-targeted nanoparticles (**Figure 3B**). In contrast to male mice, female mice demonstrated uptake by more cells as well as different clearance rates across the C’Dot groups, as evidenced by different slopes. The higher uptake of all three C’Dot types in female mice over the first 30 minutes as compared to males is consistent with our initial findings on C’Dots (42). Here, RGD-C’Dots were taken up the least and cleared fastest, PEG-C’Dots were taken up the most and cleared slower than RGD-C’Dots, while initial TAT-C’Dot uptake was intermediate, followed by continuous uptake depicted by a positive slope (**Figure 3C**). The positive slope in female mice with TAT-C’Dots is unique across all nanoparticles studied in both sexes for the short incubation time experiments (5 min.). It suggests uptake by more cells over time, and this pattern of uptake is consistent with a cell-membrane penetrating nanoparticle that evades the endocytic uptake and recycling.

The long incubation (45 min.) study further highlighted the role of nanoparticle surface functionalization in uptake and clearance. Looking at the data (**Figure 3D-E**), it is most apparent that both male and female mice had substantially more uptake of TAT-C’Dots in osteocytes than the other two C’Dot type. The TAT-C’Dots were also retained by more cells over the study period of 2.5 hours. For PEG and RGD surface functionalized C’Dots, females again showed greater uptake of both over time as compared to males. In male mice, C’Dots were fully cleared by about 1.5 hours after injection, while females retained C’Dot signal for the entire 2.5-hour observation period (**Figure 3D-E**). The data are consistent with rapid endocytic clearance of PEG- and RGD-surface functionalized C’Dots while TAT-C’Dots are able to penetrate the cell membrane and evade endocytic cycling.

To better understand the uptake process C’Dots, we examined the subcellular localization of the nanoparticles at the first imaging time point for both the short and long incubation studies. Example images of diffuse and discrete cellular signals are shown in **Figure 4A**). During the short incubation, no differences in subcellular localization were observed between the C’Dot groups for male or female mice (**Figure 4B-C**). In contrast, after the long incubation, TAT-C’Dots displayed lower levels of discrete subcellular localization in male mice compared to the other C’Dot types (**Figure 4D**). Cells treated with TAT-C’Dots exhibited a more diffuse signal, with the nanoparticles spreading evenly throughout the cytosol. This supports the conclusion that TAT-C’Dots evade endosomal regulation, affirming their role as a positive control for bypassing endocytosis in osteocytes *in vivo*. Surprisingly, females displayed no differences in subcellular localization for any C’Dot functionalization after the long incubation.

**Fig. 4.**
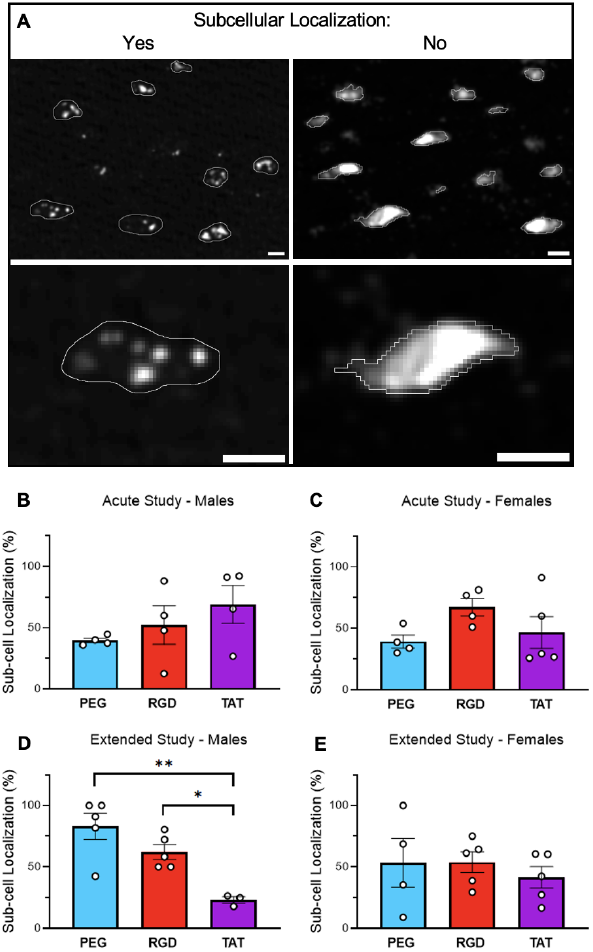
Subcellular localization of C’Dots within osteocytes in vivo is modulated by C’Dot surface functionalization in males. C’Dot signal within osteocytes can be discrete and punctate or diffuse and saturated (A, left and right, respectively). Quantification of the percentage of cells with discrete subcellular signal at the first short incubation and first long incubation experimental imaging timepoints with different C’Dot surface functionalization (B-E). Student’s T-Tests compare each of the groups within each plot (*p≤0.05, Error = SEM).

### 2.3 Osteocyte nanoparticle uptake and clearance is modulated with a cholesterol depleting agent

After establishing baseline nanoparticle behavior across different C’Dot surface functionalizations, we next examined the effect of inhibiting endocytosis on C’Dot uptake and clearance. To do this, we applied locally injected small-molecule inhibitors to disrupt osteocyte nanoparticle uptake *in vivo*, beginning with M*β*CD, a cholesterol-depleting agent. A 10 *µ*L bolus of 30 mM M*β*CD was pre-incubated in the mouse paw prior to C’Dot injection (**Supplemental Figure 2**). This procedure was carried out for both short (5 min) and long (45 min) C’Dot incubation studies.

In the short incubation study, representative images qualitatively showed the effects of M*β*CD in reducing initial C’Dot uptake across both sexes and nanoparticle types (**Supplemental Figure 3**). Quantitatively, M*β*CD lowered C’Dot uptake compared to controls in both male and female mice for most C’Dot groups (**Figure 5**), only RGD-C’Dots shown), with TAT-C’Dots in males being the singular exception (**Supplemental Figure 4**). However, M*β*CD had no effect on the rate of clearance in most groups. These findings indicate that M*β*CD inhibits initial osteocyte uptake of nanoparticles. Additionally, M*β*CD significantly increased discrete subcellular localization for PEG- and RGD-C’Dot groups in male mice (**Figure 5C**), PEG data not shown). These results further support the conclusion that cholesterol depletion influences acute nanoparticle uptake and intracellular distribution in osteocytes *in vivo*.

**Fig. 5.**
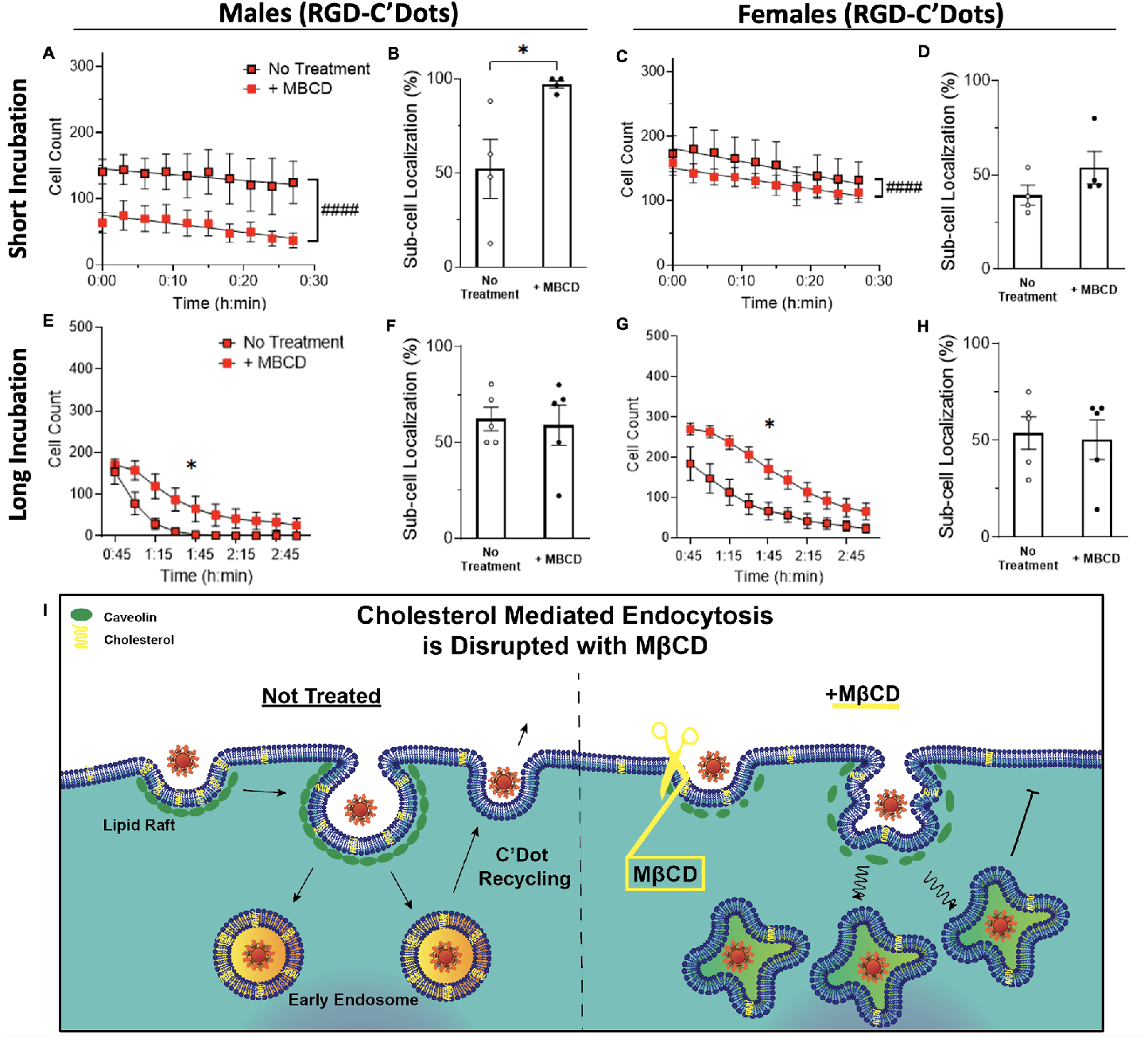
Osteocyte uptake and subcellular localization of RGD-C’Dots is modulated with the cholesterol depletor MBCD. Lipid raft mediated endocytosis is a non-sensitive uptake pathway that is disrupted with application of MBCD (A). Short incubation experiments highlight changes between non-treated and MBCD treated groups over time via linear regressions of cell count (B, D) and t-tests of subcellular localization (C,E). Long incubation experiments also show changes over time between the two-groups using a 2-Way ANOVA with repeated measures. Significant differences in the y-intercept are shown as #p≤0.05, ##p≤0.01, ###p≤0.001, ####p≤0.0001 while other comparisons show *p≤0.05, **p≤0.01, ***p≤0.001, ****p≤0.0001. Error = SEM. The response of other C’Dot surface functionalized groups to M*β*CD preincubation is shown in Supplemental Figures 4 and 5.

The effects of M*β*CD remained widespread in the long incubation study. M*β*CD pre-incubation increased osteocyte uptake of receptor-targeted RGD-C’Dots in both males and females (**Figure 5**, 2-Way ANOVAs), and a similar pattern was observed for untargeted PEG-C’Dots (**Supplemental Figure 5**). Interestingly, the opposite effect occurred in the TAT-C’Dot groups, where M*β*CD reduced osteocyte uptake of C’Dots in both sexes (**Supplemental Figure 5**). Regression analyses confirmed that M*β*CD affected the clearance rates of nearly all groups except for male PEG-C’Dots (**Supplemental Table 1, Supplemental Figure 6**).

**Fig. 6.**
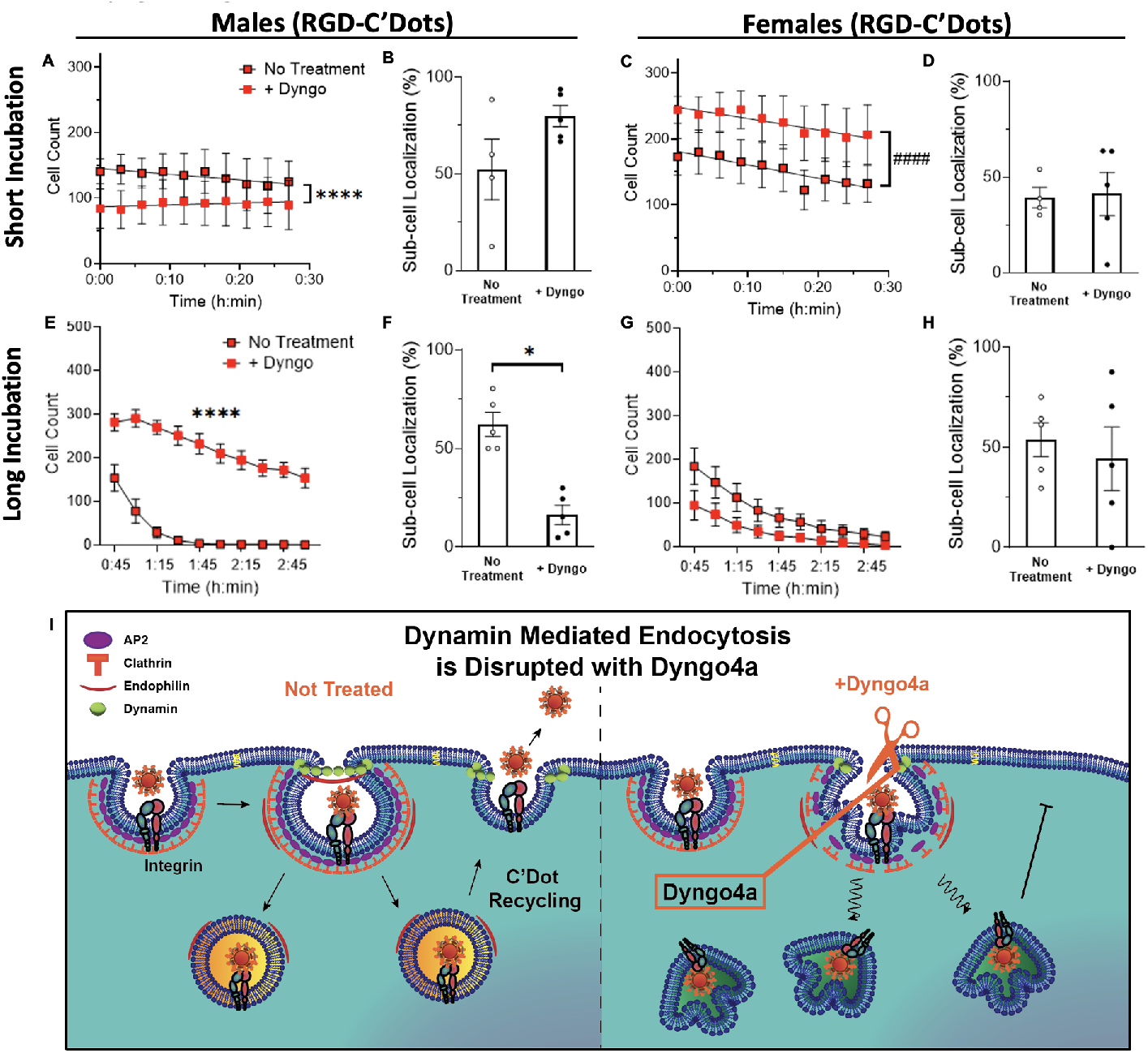
Osteocyte uptake and subcellular localization of RGD-C’Dots is modulated with the dynamin disruptor Dyngo4a. Receptor–mediated endocytosis is a sensitive uptake pathway that occurs after a receptor has been bound and is disrupted with application of Dyngo4a (A). Short incubation experiments highlight changes non-treated and Dyngo4a treated groups over time via linear regressions of cell count (B, D) and t-tests of subcellular localization (C,E). Long incubation experiments also show changes over time between the two-groups using a 2-Way ANOVA with repeated measures. Significant differences in the y-intercept are shown as #p≤0.05, ##p≤0.01, ###p≤0.001, ####p≤0.0001 while other comparisons show *p≤0.05, **p≤0.01, ***p≤0.001, ****p≤0.0001. Error = SEM. The response of other C’Dot surface functionalized groups to Dyngo4a preincubation is shown in Supplemental Figures 4 and 5.

Overall, these results demonstrate that M*β*CD has a broad impact on osteocyte endocytic dynamics, with effects that depend on surface functionalization of the nanoparticles. The data suggest that cholesterol-mediated pathways play a significant role in both the uptake and clearance of C’Dots, with distinct responses observed depending on sex and nanoparticle type.

### 2.4 Dynamin disruption impacts receptor-mediated uptake pathways in a sexually dimorphic manner

To investigate a different endocytosis pathway, we used Dyngo4a, a dynamin inhibitor, to disrupt nanoparticle uptake in osteocytes *in vivo*. Dyngo4a specifically targets receptor-mediated endocytosis, and we hypothesized it would have a more targeted effect compared to the broad activity of M*β*CD. We subcutaneously injected 10 *µ*L of 30 *µ*M Dyngo4a above the mouse metatarsal and incubated the drug for 30 minutes. We then injected C’Dots and incubated for short (5 min) and long (45 min)studies followed by time course imaging, as described previously.

In the short incubation study, pre-incubation with Dyngo4a reduced initial uptake of C’Dots in male RGD-C’Dot and female PEG-C’Dot groups. In contrast, other groups—including RGD- and TAT-C’Dots in females, and PEG- and TAT-C’Dots in males—exhibited increased C’Dot uptake compared to controls (**Supplemental Figure 4**). These findings suggest that Dyngo4a affects acute C’Dot uptake and clearance in osteocytes. Interestingly, the male RGD-C’Dot group with Dyngo4a pre-incubation displayed a positive slope, indicating a low initial uptake followed by increasing uptake over time (**Figure 6A**). However, Dyngo4a had no effect on C’Dot subcellular localization in the short study as compared to the vehicle control (**Figure 6B,D**).

In the long incubation study, dynamin inhibition increased uptake of RGD-C’Dots in male mice over time, while no changes were observed in other C’Dot types in either males or females (**Supplemental Figure 5**). In untreated males, the RGD-C’Dot signal declined after the first hour of imaging, but Dyngo4a pre-incubation caused the signal to increase and remain elevated for the duration of the experiment (**Figure 6**). This suggests a strong link between dynamin activity and the dynamics of receptor-targeted C’Dot uptake in males. In contrast, females treated with Dyngo4a showed no change in uptake or clearance of C’Dots, suggesting a sexually dimorphic response to dynamin inhibition (**Supplemental Figure 7**). A mixed model analysis confirmed that the effect of Dyngo4a on the male RGD-C’Dot group (**Supplemental Figure 8**). Regression analyses showed that the control and Dyngo4a-treated group slopes were different for RGD-C’Dots in both males and females, underscoring the role of dynamin in regulating receptor-targeted nanoparticle uptake and clearance (Supplemental Table 1). Additionally, Dyngo4a pre-incubation dramatically reduced the percentage of cells with subcellular localization to 25% in the male RGD-C’Dot group (**Figure 6**), while no other groups showed changes. These results indicate that Dyngo4a disrupts the normal uptake pathway for RGD-C’Dots in males, causing a dysregulation of receptor-mediated endocytosis.

In summary, our findings demonstrate that dynamin inhibition by Dyngo4a strongly affects the extended dynamics of receptor-targeting C’Dot uptake in males. This response is attenuated in females, highlighting a sexually dimorphic response to endocytic pathway disruption.

### 2.5 Fit of regression analysis depends on C’Dot functionalization, drug treatment, and sex

In the above results, we used linear regression to describe the C’Dot uptake and clearance data, providing a straightforward mathematical approach to quantify changes in slope and y-intercepts. While effective, this method may oversimplify the biological dynamics. This is of particular concern when signal decays in a non-linear fashion, as observed in several extended C’Dot incubation studies. To better capture the complexity of these dynamics, we applied one-phase decay (OPD) regression, which accounts for non-linear decay and can more accurately represent signal clearance profiles.

We employed OPD regression to assess the clearance kinetics (linear or decay) for each C’Dot group in the extended study (**Supplemental Table 1**). For each dataset, we tested whether a linear or OPD fit better represented the observed data. Untargeted PEG-C’Dots were generally better represented by OPD, except in M*β*CD-treated females. This decay curve suggests rapid clearance of PEG-C’Dots from osteocytes *in vivo* (**Supplemental Table 2**). In contrast, integrin-targeted RGD-C’Dots showed more variability, with a linear fit being more appropriate for males treated with Dyngo4a and females treated with M*β*CD. Interestingly, the membrane-penetrating TAT-C’Dots were consistently best represented by a linear fit across all drug treatment groups. This suggests more stable and regular retention of TAT-C’Dots within the cells over time, further supporting their ability to evade endocytic clearance mechanisms.

### 2.6 Combination plots highlight the major takeaways from endocytic modulation studies

The overall kinetics of C’Dots can be visualized by combining data points from both the short and long studies into a single, merged plot for each group (**Supplemental Figure 9**). We traced the nanoparticle uptake and clearance across both incubation studies to assess temporal changes in C’Dot kinetics and to understand the influence of drug pre-incubation on C’Dot clearance over time. Starting in the groups with no drug treatment, the uptake of PEG- and RGD-C’Dots follows an expected pattern of continuous decay for both males and females (**Supplemental Figure 9**). This trend depicts rapid clearance of C’Dot signal from osteocytes. Importantly, TAT-C’Dots have a much higher level of cellular uptake in the long incubation study for both males and females, again highlighting TAT-C’Dots as membrane-penetrating nanoparticles, evading endocytic pathways and accumulating in the cytoplasm over time.

Pre-incubation with M*β*CD resulted in an increase in extended signal for nearly all groups compared to the short study results (**Supplemental Figure 9**). T-tests confirm the change in level of C’Dot uptake between short and long timepoints. Conversely, pre-incubation with Dyngo4a resulted in an increase in extended signal for only one group, male RGD-C’Dots (**Supplemental Figure 9**). There was no impact of Dyngo4a on either PEG-or TAT-C’Dot groups compared to their controls, confirming the impact of Dyngo4a on receptor-targeted C’Dots alone.

The combined results act as an overview of our findings and clearly display the sexual dimorphism observed throughout our analyses. Three major takeaways are seen when male and female results are juxtaposed across the combined results. First, in the undrugged groups, nanoparticles are cleared more rapidly in males than in females. This is true for both untargeted and receptor-targeted C’Dots. Second, M*β*CD application increased C’Dot uptake across both male and female groups, indicating an important role for cholesterol in nanoparticle uptake across sexes. Third, Dyngo4a application does not impact females, while uptake of C’Dots is greatly prolonged in males, specifically for receptor-targeted RGD-C’Dots. A mixed model analysis confirmed sexual dimorphism across drug and C’Dot groups (**Supplemental Figure 8**).

## 3. Discussion

The concept of cellular endocytosis emerged in the late 19th century, following the first observations of phagocytosis (48, 49). Since then, endocytosis has been recognized as a key mediator of cellular responses to the extracellular environment. Endocytic mechanisms regulate the spatiotemporal composition of the plasma membrane, primarily by controlling the location, number, and availability of transmembrane proteins (50). For example, neuronal excitability depends on the precise endocytic regulation of synaptic receptors (51). Similarly, proper mechanosensitivity in fibroblasts and neutrophils requires the internalization and trafficking of integrin proteins (13, 52).

In this study, we report novel findings demonstrating that endocytic dynamics can be observed and modulated in realtime at the cellular level in live mice. What’s more, we have performed this work in one of the most challenging intravital imaging environments. Our results establish the feasibility of visualizing uptake and clearance pathways in bone cells using fluorescent nanoparticles and multiphoton microscopy (**Supplemental Videos 1, 2**). Ultrasmall fluorescent core-shell silica nanoparticles are introduced here as a tool to assess the uptake, retention, and clearance dynamics of osteocytes *in vivo*, both in the presence and absence of endocytosis-disrupting drugs. This approach represents a significant technical advance in studying live sub-cellular dynamics *in vivo* and can be easily adapted to other tissues or cell types.

Osteocytes convert tissue level mechanical cues into paracrine and endocrine signals for other cells, playing a crucial role in maintaining bone homeostasis. However, osteocytes perform their activities within a mineralized bone matrix. The central role of osteocytes in bone biology has been recognized over the past 30 years, but studying their cellular and molecular processes in live tissue remains technically challenging and rare (37, 39, 53, 54). As a result, many fundamental cellular processes (i.e., endocytosis) are still not well understood in osteocytes.

The data presented here are the first examples of using intravital multiphoton imaging to observe endocytic dynamics in osteocytes *in vivo*. They offer evidence that osteocytes use at least two pathways of endocytosis for internalizing locally delivered nanoparticles *in vivo*. We also report important differences in endocytic dynamics between male and female mice that were previously unknown. With this work, we have established a flexible platform for interrogating endocytosis dynamics *in vivo* that leverages nanoparticle surface functionalization to target specific endocytic uptake mechanisms and pharmacological agents to disrupt specific pathway components. Our contributions here represent a major leap forward in understanding and technical ability for studying endocytosis generally, and bone biology specifically.

The effects of endocytosis-inhibiting drugs *in vivo* remain largely uncharacterized (55). Our results demonstrate the effects of plasma membrane cholesterol modulation (i.e., M*β*CD treatment) on nanoparticle uptake. We saw responses across nanoparticle functionalizations and mouse sex. These findings indicate that cholesterol and lipid raft modulation can broadly influence cellular uptake (**Figure 5, Supplemental Figures 3, 5**). This result is corroborated by other researchers (56, 57). In one study, endocytosis of activated receptors was shown to limit the mechanosensitive response in bone tissue (58). Disruption of caveolin proteins in MC3T3-E1 pre-osteoblasts *in vitro* led to increased calcium signaling and enhanced mineralization in response to ATP stimulation, highlighting the potential of targeting endocytic pathways to enhance mechanosensitivity in bone (58). This approach could be particularly beneficial for aging and osteoporotic patients, where strengthening the mechanosensitive response is a key therapeutic goal. Our findings indicate that membrane cholesterol and cholesterol-mediated endocytosis are also broadly utilized in osteocytes. Targeting these pathways in osteocytes specifically could offer a novel therapeutic strategy for modulating bone’s response to mechanical stimulation, making it a promising area for future research in bone health and disease.

We noted additional patterns in the drug modulated uptake kinetics of nanoparticles over time. Several drug-treated groups displayed a strong rebound in C’Dot uptake between the short and long incubation studies (**Supplemental Figure 9**). This suggests that the small molecule treatments initially inhibited normal C’Dot uptake, before allowing later C’Dot accumulation in the cells. *in vitro* studies indicate that the effects of M*β*CD-induced membrane cholesterol depletion can be long-lasting, persisting for at least 12 hours (59, 60). In contrast, *in vitro* studies with Dyngo4a report a more transient effect, with dynamin-mediated endocytosis recovering within 30 minutes after drug washout (19). It is important to note that mice were allowed normal cage activity during the extended (45 minute) C’Dot incubation, whereas they remained under anesthesia during the short (5 minutes) C’Dot incubation. Previous models and *in vivo* studies suggest that mechanical loading enhances oscillatory fluid flow and tracer entry into the osteocyte extracellular microenvironment (61, 62). Increased fluid flow may partially explain the differences observed between the short and long incubation studies, however the combined plots of untreated groups follows an expected progression of clearance (**Supplemental Figure 9**). Additionally, other *in vivo* endocytosis experiments have reported increased uptake of exogenously applied dextran over time, as endocytic trafficking and vesicle fusion progress (63). This could be another explanation for the increased C’Dot uptake in some of our long incubation time points. However, in that case we would expect a pattern of delayed uptake across all groups, instead of only drug treated groups. Future research will need to investigate the active duration of these small molecule endocytosis inhibitors *in vivo* to determine whether they can be used to modulate cell metabolism in a transient manner.

C’Dot functionalization provided unique insight into osteocyte endocytic pathways. All three C’Dot types displayed distinct uptake and clearance profiles, especially visible in the long incubation study. Cell membrane penetrating TAT-C’Dots maintained high levels of cell uptake over time, aligning with the hypothesis that these C’Dots evade endocytic trafficking and localize indelibly to the cytoplasm (**Figure 3, Supplemental Figure 10**). Untargeted PEG-C’Dot uptake was modulated by cholesterol depletion but not by dynamin disruption, which suggests that these nanoparticles are being taken up by non-specific endocytic pathways. Finally, integrin-targeted RGD-C’Dots were highly impacted by Dyngo preincubation, but only in males (**Figure 6**). This suggests that dynamin GTPase activity is critical for rapid clearance and subcellular localization of integrin targeted nanoparticles in male but not female mice. This information is especially thought-provoking in light of the critical role of integrins and other transmembrane receptors in osteocyte mechanosensitivity both *in vitro* and *in vivo* (64–69).

Although we did not anticipate sex differences in this cellular-level study of endocytosis, our results reveal a clear pattern of sexual dimorphism. At baseline, the clearance of both untargeted and integrin-targeted C’Dots varied between male and female mice, with female mice retaining C’Dot signal for longer (**Supplemental Figure 8**). Drug treatments also had sex-specific effects. Cholesterol modulation increased uptake of PEG- and RGD-C’Dots in the long study in both sexes. However, dynamin inhibition impacted the sexes in opposite ways. Linear regressions of untargeted and integrin-targeted C’Dots in the long study showed that females displayed a modest decrease in C’Dot uptake after Dyngo4a pre-incubation. In contrast, males displayed sizeable increase in uptake, particularly with the RGD-C’Dots (**Supplemental Figure 6**). Given the general ubiquity of endocytic pathways, it is intriguing that the clearance dynamics of C’Dots exhibited sexual dimorphism. The minimal response to Dyngo4a pre-incubation by females suggests that the canonical clathrin-mediated endocytic pathway may play a lesser role in nanoparticle metabolism in female osteocytes. Beyond C’Dot uptake, the subcellular localization of C’Dots also differed between the sexes (**Figure 4**). Notably, females did not exhibit changes in subcellular localization in response to nanoparticle functionalization or drug treatment, unlike males. Although our current imaging resolution does not allow us to directly visualize endosomes or lysosomes, the observed punctate signal may indicate a more organized or compact uptake process in males compared to the diffuse signal seen in females. This suggests that cholesterol and dynamin inhibition may disrupt normal vesicular packaging of nanoparticles at different time points in males. Overall, our findings highlight a previously unrecognized sexual dimorphism in endocytic mechanisms. Future research should investigate whether differential expression of endocytic components, such as dynamin, exists between males and females at the cellular level.

The findings reported here provide the first quantification of a novel platform for investigating live endocytosis dynamics *in vivo*. Our study lays the groundwork for future research exploring how endocytic disruption affects the availability of mechanosensitive proteins and, consequently, osteocyte mechanosensitivity. Ongoing studies in our lab will address this by examining the downstream mechanosensitive responses of osteocytes to endocytic inhibitors *in vivo*. If the localization and availability of mechanosensitive proteins can be pharmacologically controlled, disrupting specific endocytic pathways may offer a new strategy for modulating cellular responses to mechanical stimulation *in vivo*. Furthermore, the unexpected observation of sexual dimorphism in endocytic pathway utilization is a compelling finding that warrants further investigation within the scientific community.

## 4. Methods

### 4.1 Animals

Male and female skeletally mature young adult C57Bl/6J mice (16-20 weeks old) were used for all nanoparticle uptake and clearance experiments (Jackson Laboratory, Bar Harbor, ME). Intracellular co-localization experiments utilized an established mouse line with osteocyte-targeted expression of the genetically encoded calcium indicator (GECI) GCaMP6f (46). These mice were bred in house by crossing Ai95-D mice [B6J.Cg-Gt(ROSA)26Sortm95.1(CAGGCaMP6f)Hze/MwarJ; JAX Labs] with DMP1/Cre mice [B6N.FVB-Tg(Dmp1-cre)1Jqfe/BwdJ; JAX Labs], which have Cre recombinase driven by the DMP1 promoter, a gene predominantly expressed in osteocytes (70). All procedures were approved by the Institutional Animal Care and Use Committees of Cornell University.

### 4.2 Synthesis of ultrasmall PEGylated, cRGD, and TAT functionalized Core-Shell Silica Nanoparticles

Ultrasmall fluorescent core-shell silica nanoparticles (C’Dots) with covalently encapsulated Cy5 dye (from Cy5-maleimide (Lumiprobe) with net positive charge conjugated to (3-mercaptopropyl)trimethoxysilane (MPTMS; Sigma-Aldrich) in a silica core (from TMOS; Sigma-Aldrich) and poly(ethylene glycol) (PEG) shell (from methoxy-PEG(6-9)-silane (500 g/mol); Gelest) were synthesized in water as previously described (43, 71). Three varieties of C’Dots were synthesized: 1) Untargeted control C’Dots (PEG-Cy5-C’Dots) - referred to as PEG-C’Dots. 2) Integrin targeting C’Dots surface functionalized with integrin-targeting cyclic(arginine-glycine-aspartic acid-D, tyrosine-cysteine) peptide, (c(RGDyC)-PEG-Cy5-C’Dots) - referred to as RGD-C’Dots. RGD-C’Dots were functionalized by adding cRGDyC-PEG-silane to the particle core in the PEGylation step as described in detail elsewhere (43, 44). 3) C’Dots capable of penetrating membranes, including endosomal escape, were surface functionalized with human immunodeficiency virus (HIV) TAT peptides (TAT-PEG-Cy5-C’Dots) - referred to as TAT-C’Dots. TAT-C’Dots were prepared similar to previously reported methods (72), but using an improved click chemistry approach. The TAT peptide sequence used was 5-azido-pentanoyl – RKKRRQRRR-NH2 (Biosynth). The terminal azido group of the peptide was clicked onto the C’Dot surface functionalized with dibenzocyclooctyne (DBCO) via azide-alkyne cycloaddition. The DBCO functionalized C’Dots were prepared using post-PEGylation surface modification by insertion (PPSMI) method as described in detail elsewhere (73). To that end, small functional aminopropyl-trimethoxysilane (APTMS) was first inserted between PEGs on the silica surface. Resulting NH2-C’ dots were then reacted with DBCO-PEG4-Nhydroxysuccinimidyl ester (DBCO-PEG4-NHS ester, Santa Cruz Biotechnology) yielding DBCO-C’Dots. From particle characterization efforts employing a combination of fluorescence correlation spectroscopy (FCS) and UV-Vis absorbance data, with methods described elsewhere (73), PEG-, RGD-, and TAT-C’Dots exhibited hydrodynamic radii of 5.7, 5.5, and 6.0nm, respectively, and number of dyes per C’Dot of 2.0, 2.1, and 2.0, respectively (**Supplemental Figure 12**. Furthermore, RGD-C’Dots were estimated to have 20 cRGDyC ligands per particle and TAT-C’Dots were estimated to have 10 TAT (5-azido-pentanoyl – RKKRRQRRRNH2) peptides per particle.

### 4.3 Endocytosis Modulating Drugs

The small molecule Dyngo4a (Abcam) was used as a potent Dynasore analog to evaluate the impact of dynamin-mediated endocytosis inhibition on osteocyte C’Dot uptake and metabolism *in vivo*. The molecule disrupts dynamin GTPase activity and its role in pinching off budding endocytic vesicles (19). Dyngo4a was diluted from a stock solution in DMSO to the working concentration of 30*µ*M in 1x DPBS. Dyngo4a working solution was kept at -20°C and was brought to room temperature before use. Water-soluble oligosaccharide methyl-*β*-cyclodextrin (M*β*CD, Sigma) binds to and removes cholesterol from the plasma membrane and is used to evaluate the impact of cholesterol-mediated lipid raft-based endocytosis inhibition (22). M*β*CD was diluted from a powder to its working concentration of 10mM in 1x DPBS. M*β*CD was kept at 4°C and brought to room temperature before use.

### 4.4 Drug Injection and Incubation

Mice were anesthetized with 2-3% isoflurane mixed with medical grade air in an induction chamber for 3 minutes before injection. Restraint of the mice was necessary to maintain reliable reproduction of the precise injection procedure. During injection, mice were kept under 2% isoflurane with a nose cone. A 10*µ*L bolus was subcutaneously injected above the MT3 30 minutes before C’Dot injection and subsequent imaging. A local injection was used to minimize the off-target endocrine effects of a systemic injection. A 10*µ*L control bolus of 1x DPBS was used on control groups in the acute clearance study. No control bolus was used in the extended clearance study. Mice were allowed normal cage activity during the 30-minute drug pre-incubation time.

### 4.5 C’Dot Injection and Incubation

After drug or control bolus preincubation, mice were re-anesthetized for nanoparticle injection. A 10*µ*L subcutaneous injection of 10*µ*M PEG-, RGD-, or TAT-C’Dots was then administered over the third metatarsal (MT3). For the acute clearance study, C’Dots were incubated for 5 minutes and mice were kept under nose cone anesthesia before MT3 isolation and imaging (n=4-5/group/sex). For the extended clearance study, C’Dots were incubated for 45 minutes before MT3 isolation. Mice were allowed normal cage activity during the 45-minute incubation and were re-anesthetized immediately before MT3 isolation surgery (n=3-5/group/sex).

### 4.6 Metatarsal Isolation Surgery

MT3 bones were isolated as previously described (42, 45). Briefly, while mice were anesthetized, a shallow incision was made between the second and third metatarsal on the dorsal aspect of the mouse hind paw, and overlying tendons were removed. The MT3 was then functionally isolated from the rest of the paw with a stainless steel pin beneath the mid-diaphysis of the bone, leaving primary vasculature at the proximal and distal palmar aspects intact. The bone was secured in a 3-point bending configuration and submerged in room temperature 1xDPBS. Mice were continuously anesthetized with a nose cone at 1.5-2% isoflurane during all subsequent imaging.

### 4.7 Imaging

: Fluorescent signal inside the functionally isolated MT3 was visualized with multiphoton microscopy (MPM) (Bergamo II, Thorlabs) as previously described (42). Briefly, a 20x immersion objective (XLUMPLFLN, Olympus), 1090nm wavelength excitation, and a >647nm long pass filter acquisition were used for observation of Cy5 encapsulated C’Dots. Images were captured at 1024×1024 pixel density and 0.588*µ*m pixel resolution. A 35*µ*m z-stack was taken for each mouse, starting 20*µ*m beneath the bone surface, with 0.3*µ*m steps between each slice and averaging of 7 frames per slice. In the short incubation clearance study, after the 5-minute incubation, imaging occurred every 3 minutes for 30 minutes for a total of 10 stacks. During the long incubation clearance study, after the initial 45-minute incubation, imaging occurred every 15 minutes over a period of 2.5 hours, also resulting in ten stacks (experimental timelines for the short and long imaging studies are depicted in **Supplemental Figure 2**. Timelapse videos use the same pixel density and resolution, capturing an image every 30 seconds for 15 minutes. Co-localization imaging experiments with GCaMP6f mice used a second volumetric z-stack with an excitation of 920nm and 525 ± 25nm bandpass filter acquisition to collect intracellular calcium signal. Experiments looking at C’Dot dendritic signal used a 5-minute C’Dot incubation before MT3 isolation. Averaging was increased to 20 frames per slice to visualize the fine structure of osteocyte dendrites. Images showing colocalization of C’Dots and third harmonic generation (THG) signal were collected using a 3-photon system created in the Xu and Schaffer Labs at Cornell (47). Briefly, an Opera-F amplifier laser (Coherent) with an excitation of 1680nm and a 25x water-immersion objective (XLPLN25XWMP, Olympus) was used. The mouse hind paw with functionally isolated MT3 bone (as described in section 2.6) was immersed in heavy water (D2O) to reduce absorption of the excitation laser. THG signal was detected through a 558 ± 10nm bandpass filter (Semrock) by an ultra bialkali PMT (R7600-200, QE at 560 nm, 10%, Hamamatsu Photonics) while red Cy-5 C’Dot signal was detected through a 593nm long-pass filter (Semrock). A 35*µ*m z-stack was obtained with 0.3*µ*m steps between each frame. MATLAB (MathWorks) software running the ScanImage (74) module mediated image collection.

### 4.8 Image Analysis

Z-stacks for all imaging sessions of each mouse were analyzed in ImageJ (NIH) as previously described (42). Briefly, a macro was created to filter the images and to interpolate the cortical bone regions of interest across the z-stack depth. Then 3D segmentation created objects for all cell volumes over a threshold signal intensity. The threshold value was established as the mean grey value of background region of interest (ROI) plus 3x standard deviation. Segmented objects were then filtered by surface area (>1000 pixels) to ensure only full cell volumes were counted. The total number of objects with C’Dot signal was collected for each imaging time point. The mean intensity for each object was also quantified using the 3D Manager tool. Intensities were normalized to the background within each z-stack.

### 4.9 Quantification of C’Dot Subcellular Localization

The first frame from each z-stack taken after C’Dot incubations was analyzed for subcellular localization within osteocytes. PEG- and RGD-C’Dot groups were quantified under the following study conditions of: no drug, Dyno4a pre-incubation, and M*β*CD pre-incubation. Images were filtered with a Gaussian blur, thresholded, and cellular ROIs containing C’Dot signal were identified with the “Analyze Particle” tool. These ROIs were overlayed onto the original un-blurred image to assist with cell visualization. Cells were binned as having either distinct subcellular localization of C’Dots or saturated/non-distinct signal. The number of cells with subcellular localization was reported as a percentage of total cells in each frame.

### 4.10 Statistical Analyses

Clearance study analysis was performed using 2-way ANOVAs and regressions, including non-linear (one-phase decay) and linear regressions. Quantification of changes between acute and extended clearance studies used a student’s T-Test on the final acute time point and the initial extended time point. Subcellular localization analysis used student’s T-Tests between drug groups within sex and C’Dot type. Analyses were performed in GraphPad Prism version 9.0 for Mac OS X (GraphPad Software). A mixed model on extended clearance data was also run in RStudio version 2023.12.1+402 using the lme4 package.

### 4.11 Overall Study Design

Osteocytes in C57Bl/6J mouse third metatarsals were exposed to Dyngo4a and M*β*CD small molecule drugs via a local subcutaneous injection. These drugs were assessed for their ability to modulate osteocyte uptake, metabolism, and trafficking of locally delivered nanoparticles. Previously validated fluorescent nanoparticles (C’Dots) were used to observe changes in osteocyte activity *in vivo* (42). Either untargeted PEG-C’Dots, integrin-targeting RGD-C’Dots, or membrane penetrating/endosomal escaping TAT-C’Dots were injected into the mouse hind paw 30 minutes after preincubation with Dyngo4a, M*β*CD, or a control PBS bolus (**Supplemental Figure 2**. Three C’Dots types with distinct cellular uptake pathways and two broadly acting drugs were used to establish a better fundamental understanding of osteocyte nanoparticle endocytosis *in vivo*. Volumetric multiphoton near-infrared imaging of C’Dot signal over time was used to quantify drug induced changes in osteocyte C’Dot uptake and clearance. Both Dyngo4a and M*β*CD were tested across all groups: male and female, PEG-, RGD-, and TAT-C’Dots for both short and long incubation clearance studies (n=4-5/group/sex).

## ACKNOWLEDGEMENTS

Thank you to the Weill Hall CARE staff and Karly Hooper for maintaining the mouse colonies used in these experiments. Thanks to Chris Schaffer and Nozomi Nishimura for access to their 3-Photon microscope and to Samantha Bratcher for assisting with 3-Photon image collection. This work was supported by the Department of Biomedical Engineering at Cornell University.

## Supplementary Information

**Fig. 7. Figure S1:**
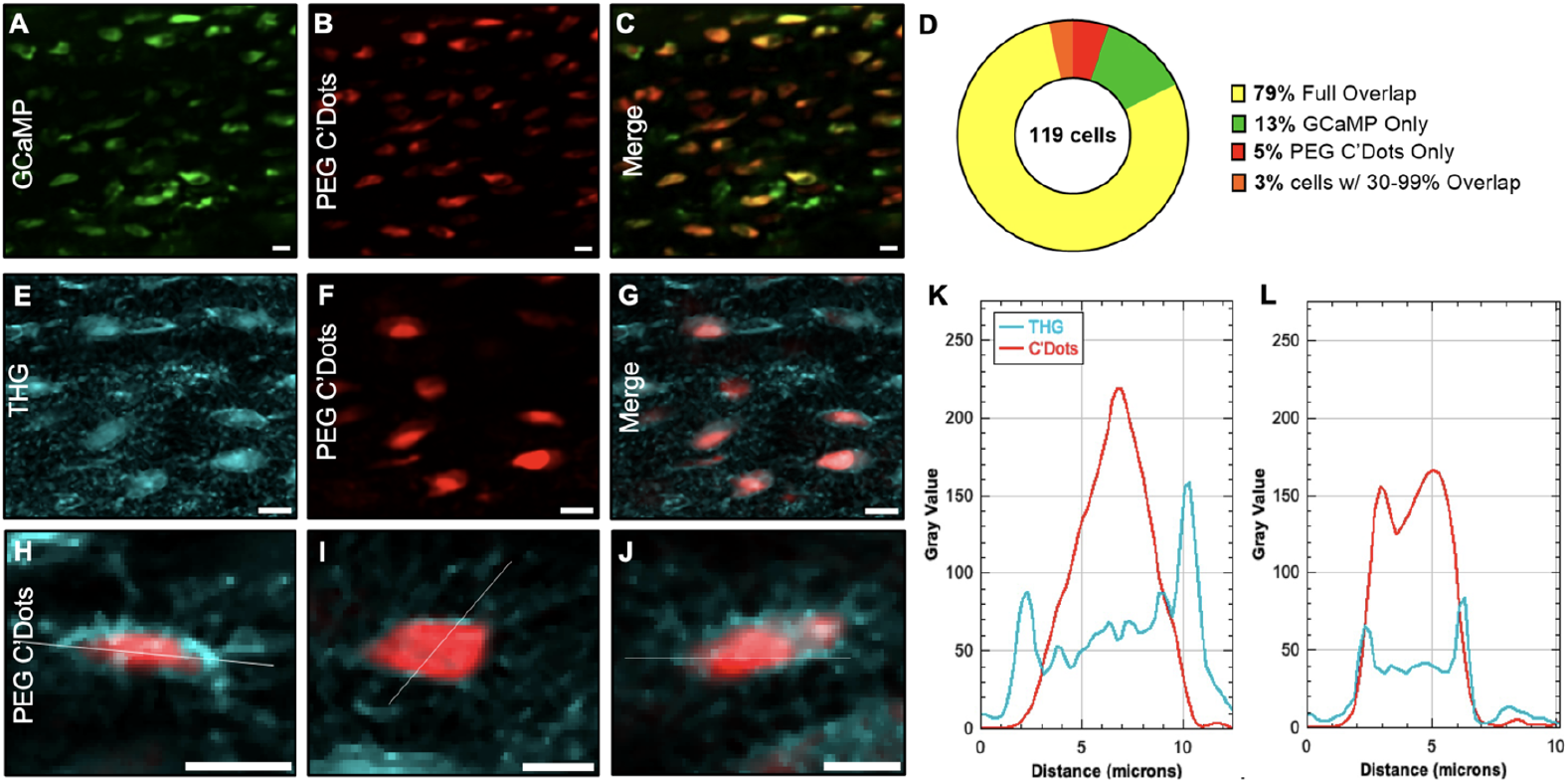
Subcutaneously injected PEG-C’Dots are internalized by osteocytes in vivo. To confirm the cellular uptake of the C’Dot nanoparticles, we performed distinct co-localization experiments with both 2-photon and 3-Photon microscopy. In the 2-Photon experiments, genetically modified mice expressing GCaMP6f signal within osteocytes were locally injected with PEG C’Dots (A-C) for a 45-minute incubation before imaging. GCaMP signal (green) was collected at 920nm excitation and C’Dot signal (red) was collected at 1090nm excitation. Co-localization analysis on the overlap of signal showed a high percentage of cells with full or partial overlap for PEG C’Dots (D). For the 3-Photon experiment, wild-type mice were locally injected with PEG-C’Dots (E-G) 45 minutes before imaging with endogenous third harmonic generation signal. Analysis of the signal intensity across a linear profile of representative cells (H-J, white line) showed that the THG signal, which highlights material interfaces (i.e. matrix/fluid boundary), was external to the PEG-C’Dots (K-L).

**Fig. 8. Figure S2:**
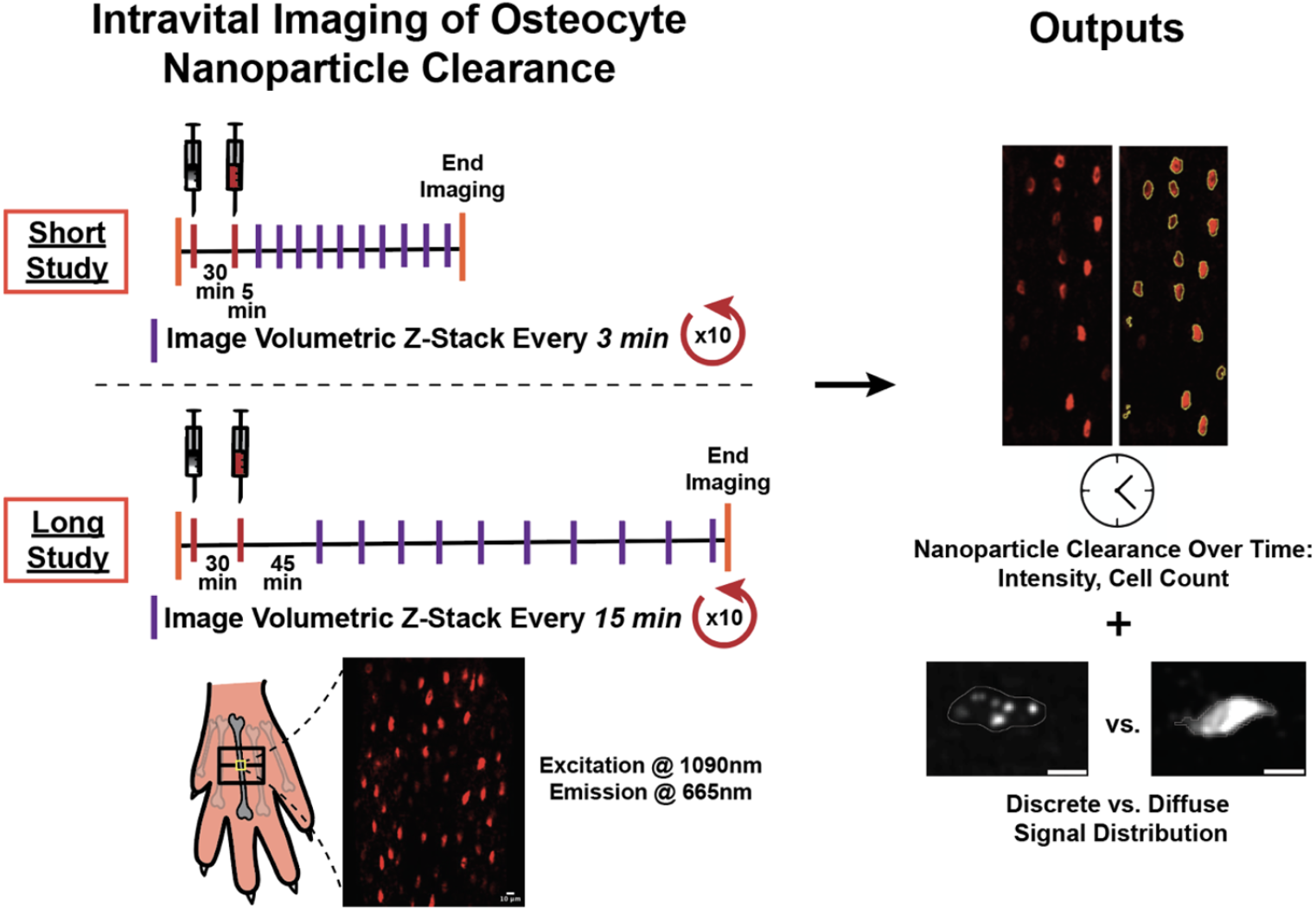
Graphical abstract of experimental design and timeline for imaging intravital of C’Dot uptake and clearance in bone with drug pre-incubation. Both endocytosis disrupting drugs and fluorescent nanoparticles were exposed to bone and osteocytes through a local subcutaneous injection above the third metatarsal (3MT) bone in a mouse hind paw. After a 30-minute pre-incubation of the drugs or control, C’Dot nanoparticles were injected and incubated before surgical MT3 isolation and imaging of cortical bone osteocytes. The short incubation experiment had a 5 min C’Dot incubation time and z-stack images were taken every 3 minutes for 30 minutes. The long incubation experiment utilized a 45-minute C’Dot incubation time and z-stack images were taken every 15 minutes for 2.5 hours. Images from both experiments were analyzed for clearance kinetics of nanoparticle signal as well as subcellular distribution of signal. Scale bars = 10*µ*m.

**Fig. 9. Figure S3:**
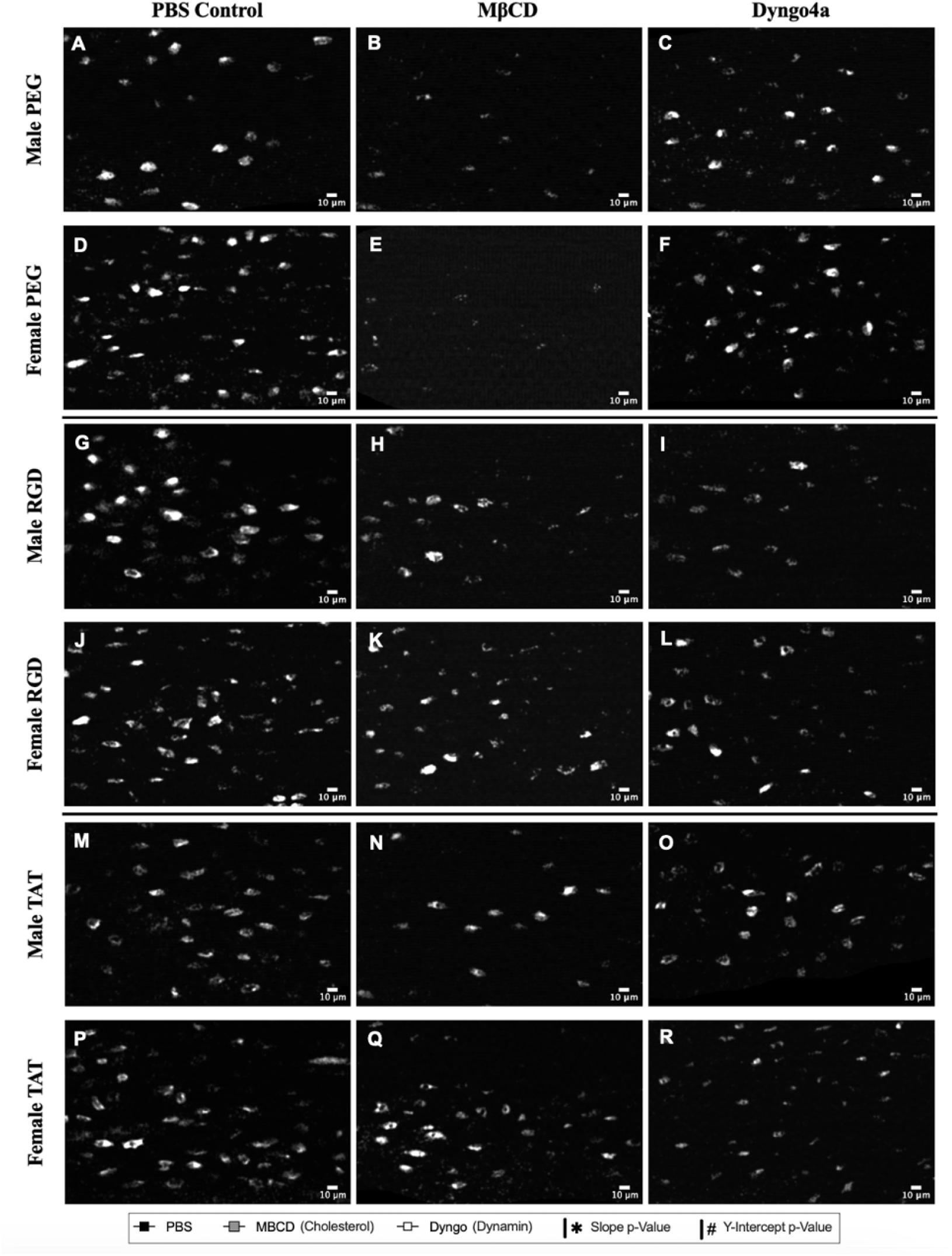
Representative single-frame images of acute C’Dot uptake into cortical bone osteocytes under different endocytic drug conditions illustrating the broad impact of M*β*CD. Images were taken immediately after a 30-minute incubation of drug or PBS control followed by a 5-minute incubation of PEG-(A-F), RGD- (G-L), or TAT- (M-R) C’Dots. Complete z-stacks from each time point were used to quantify the amount of osteocyte nanoparticle uptake over time shown in Figure 5. Scale bars = 10*µ*m.

**Fig. 10. Figure S4:**
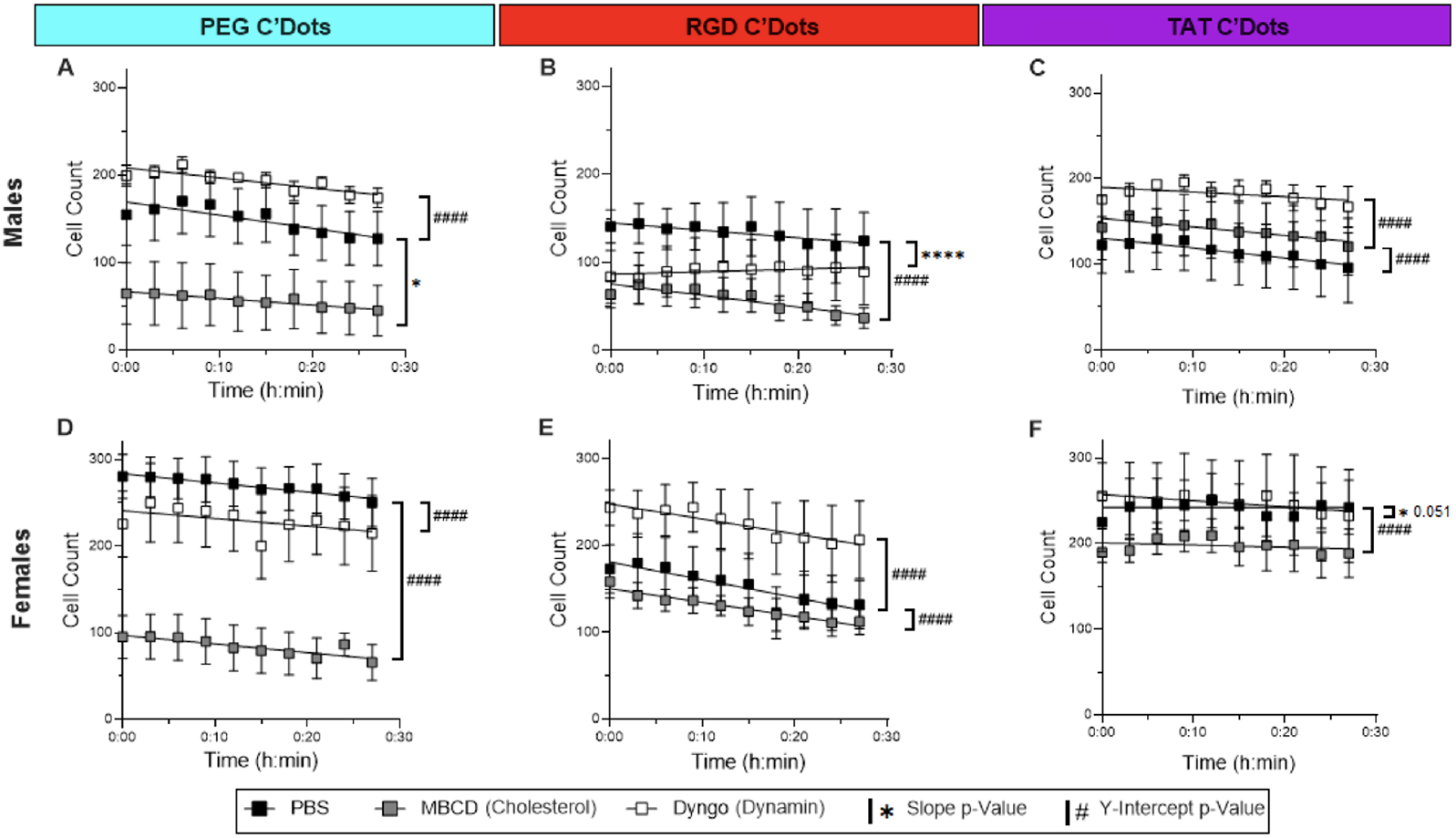
Acute C’Dot uptake and clearance in cortical bone osteocytes in vivo is modulated by endocytosis modulating drug pre-incubation. Grouped by sex and C’Dot type, simple linear regressions measure the impact of endocytic drugs on C’Dot uptake in osteocytes over time relative to PBS controls. Male groups are graphed on the top row (A-C) and female groups are graphed on the bottom row (D-F). Statistical measurements compare drug groups to PBS control within each panel. A significant difference in slope is shown as *p≤0.05, **p≤0.01, ***p≤0.001, ****p≤0.0001. A significant difference in y-intercept is shown as #p≤0.05, ##p≤0.01, ###p≤0.001, ####p≤0.0001. Error = SEM.

**Fig. 11. Figure S5:**
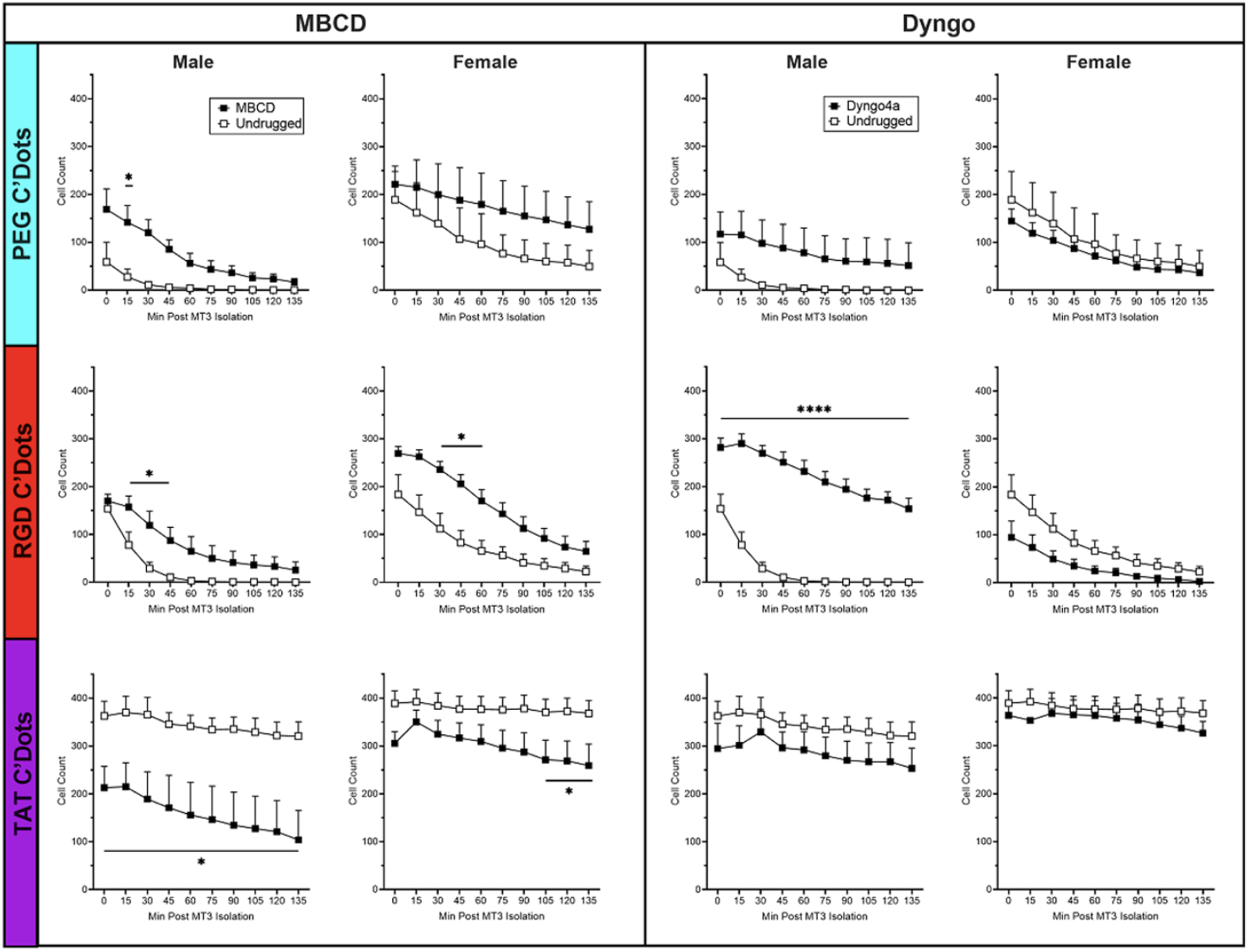
The kinetics of extended C’Dot clearance in osteocytes are broadly changed under cholesterol depletion and only changed in RGD-C’Dot Males with dynamin inhibition. After a 45-minute long C’Dot incubation, osteocyte uptake of C’Dots under different drug conditions was quantified over time (n=3-5/group). The impact of cholesterol inhibitor (MBCD) and dynamin inhibitor (Dyngo4a) compared to untreated control (black) was assessed via 2-way ANOVA. Male data was transformed (Y=log(Y+1)) due to near-zero data points before ANOVA analysis, untransformed data is shown in the graph. Stats compare the drug group to untreated control at each timepoint (*p≤0.05, **p≤0.01, ***p≤0.001, ****p≤0.0001, Error = SEM).

**Fig. 12. Supplementary Table 1:** Long incubation C’Dot clearance profiles are closely represented by linear and one phase decay regression analyses. For each C’Dot group, linear regressions (red) and One one-phase decay regressions (blue) are fitted to the data. The impact of dynamin inhibition (Dyngo) and cholesterol depletion (MBCD) on C’Dot uptake and clearance is quantified as changes in linear slope and y-intercept. Bolded sections show significant differences (p≤0.05). Graphs depicting the regression lines are shown in further supplemental data (Supplemental Figures 2, 3).

| C'Dot Treatment Group | Linear Regression (Fit – R<sup>2</sup>) | One Phase Decay (Fit – R<sup>2</sup>) | Linear Slope vs Dyngo4a | Linear Y-Intercepts vs Dyngo4a | Linear Slope vs MBCD | Linear Y-Intercepts vs MBCD |
| --- | --- | --- | --- | --- | --- | --- |
| Male PEG | 0.7679 | 0.9987 | p=0.3022 | **p<0.0001** | p=0.1226 | **p<0.0001** |
| Male RGD | 0.8685 | 0.9975 | **p=0.0018** | - | **p=0.0139** | - |
| Male TAT | 0.9138 | 0.9176 | p=0.7523 | **p<0.0001** | **p<0.0001** | - |
| Female PEG | 0.9187 | 0.9937 | p=0.0878 | **p=0.0007** | **p=0.0127** | - |
| Female RGD | 0.8976 | 0.9983 | **p=0.0067** | - | **p=0.0022** | - |
| Female TAT | 0.8511 | 0.8794 | p=0.2006 | **p<0.0001** | **p=0.0021** | - |

**Fig. 13. Supplementary Table 2:** Cholesterol depletion modulated the long incubation clearance profiles of C’Dots more broadly than dynamin disruption. A comparison test between the two regression curves defines the preferred mode of regression for each C’Dot group. The second column shows the preferred regression without any drug. The third and fourth columns highlight the impact of dynamin inhibition or cholesterol depletion on the preferred regression.

| C'Dot Treatment Group | Linear or OPD Preferred? | Linear or OPD Preferred w/Dyngo4a? | Linear or OPD Preferred w/MBCD? |
| --- | --- | --- | --- |
| Male PEG | OPD | OPD | OPD |
| Male RGD | OPD | Linear | OPD |
| Male TAT | Linear | Linear | Linear |
| Female PEG | OPD | OPD | Linear |
| Female RGD | OPD | OPD | Linear |
| Female TAT | Linear | Linear | Linear |

**Fig. 14. Figure S6:**
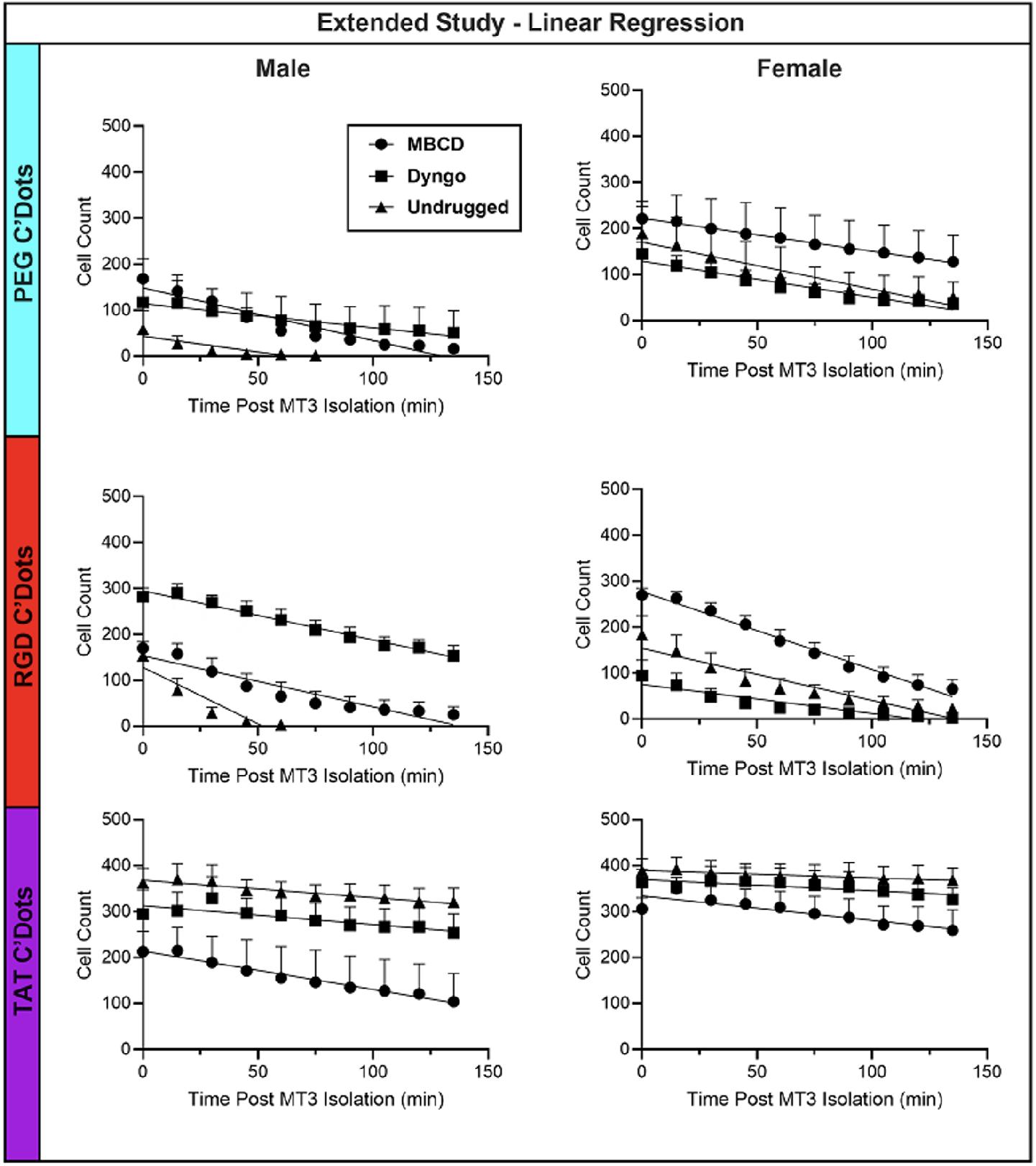
Simple linear regression lines on extended C’Dot clearance experiment groups clearly display differences in slope and y-intercept with drug pre-incubation. Data points reaching 0 (in Male PEG and RGD groups) were removed before regression analysis. N=3-5/group, Error = SEM.

**Fig. 15. Figure S7:**
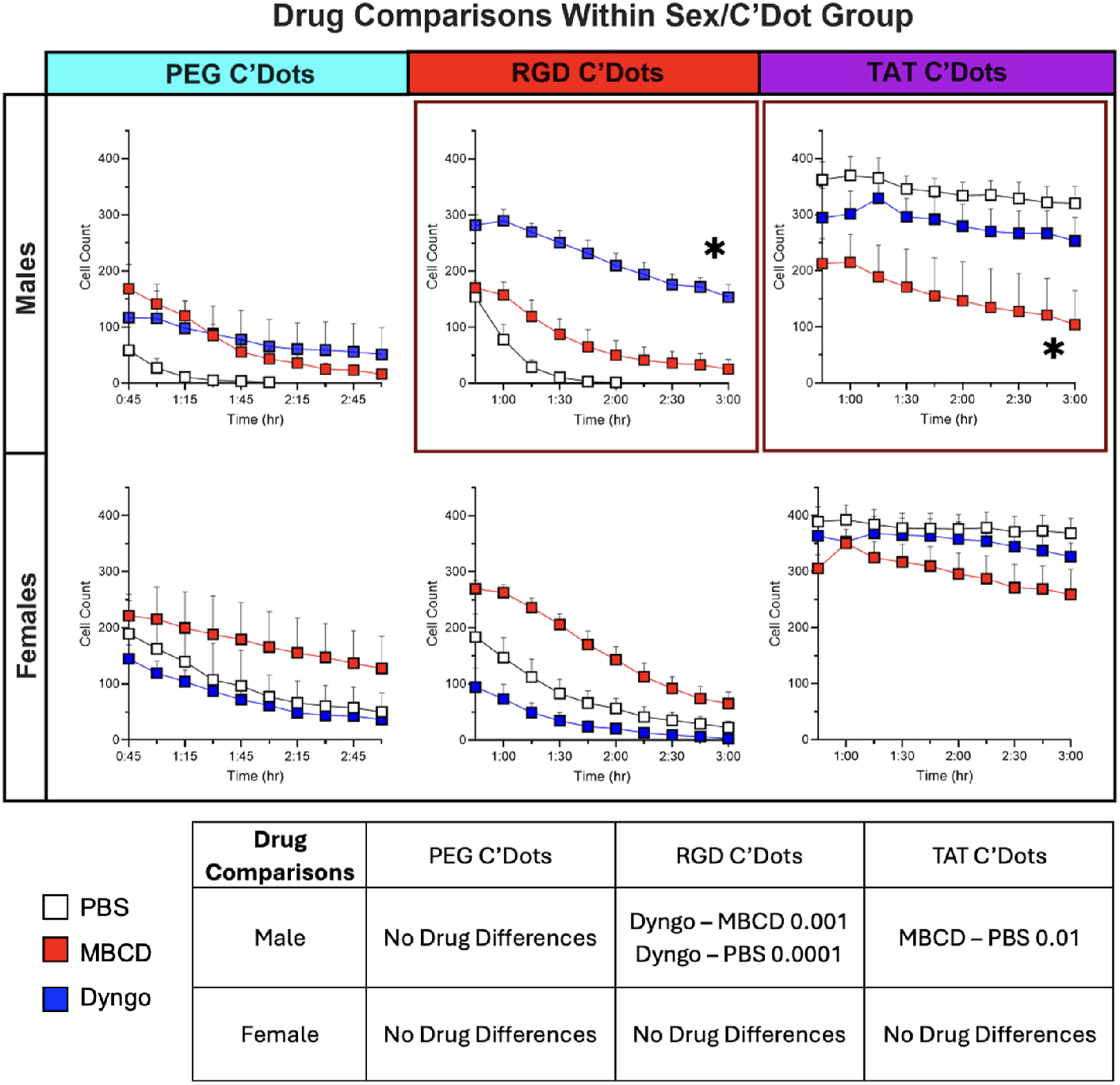
A Mixed Linear Model highlights the impact of Dyngo4a on RGD-C’Dot males and M*β*CD on TAT-C’Dot males. Plots show mixed model comparisons of drugs within C’Dot and sex groups for the long incubation clearance data. Comparisons with significant differences are boxed in red. P≤0.05, Error = ±SEM.

**Fig. 16. Figure S8:**
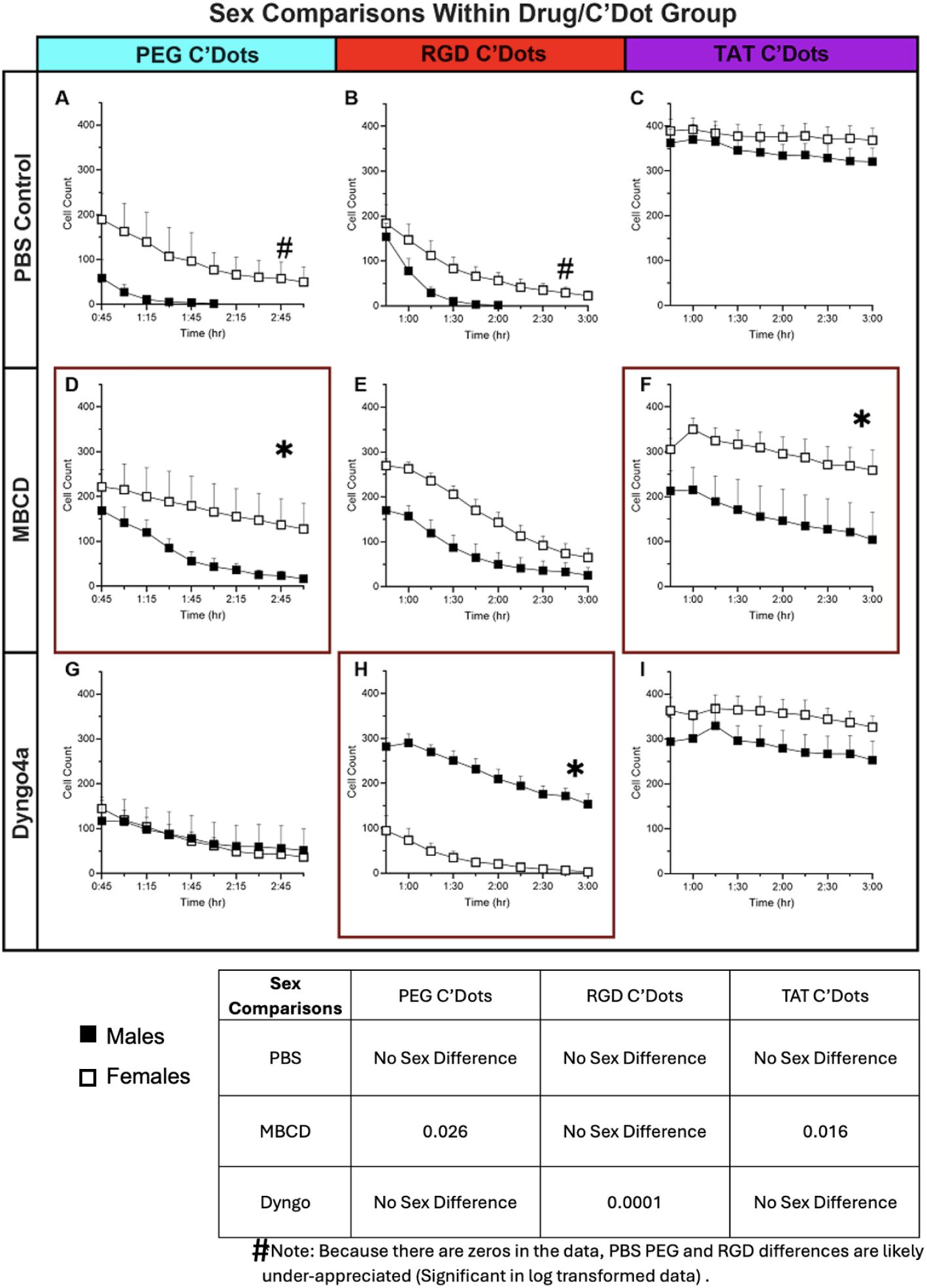
A Mixed Linear Model highlights the impact of sex on distinct drug and C’Dot groups. Plots show mixed model comparisons of sex within C’Dot and drug groups for thelong incubation clearance data. Comparisons with significant differences are boxed in red. Pound symbols denote that the untransformed data with very low or zero values may be under-appreciated in this mixed model. *P≤0.05, Error = ±SEM.

**Fig. 17. Figure S9:**
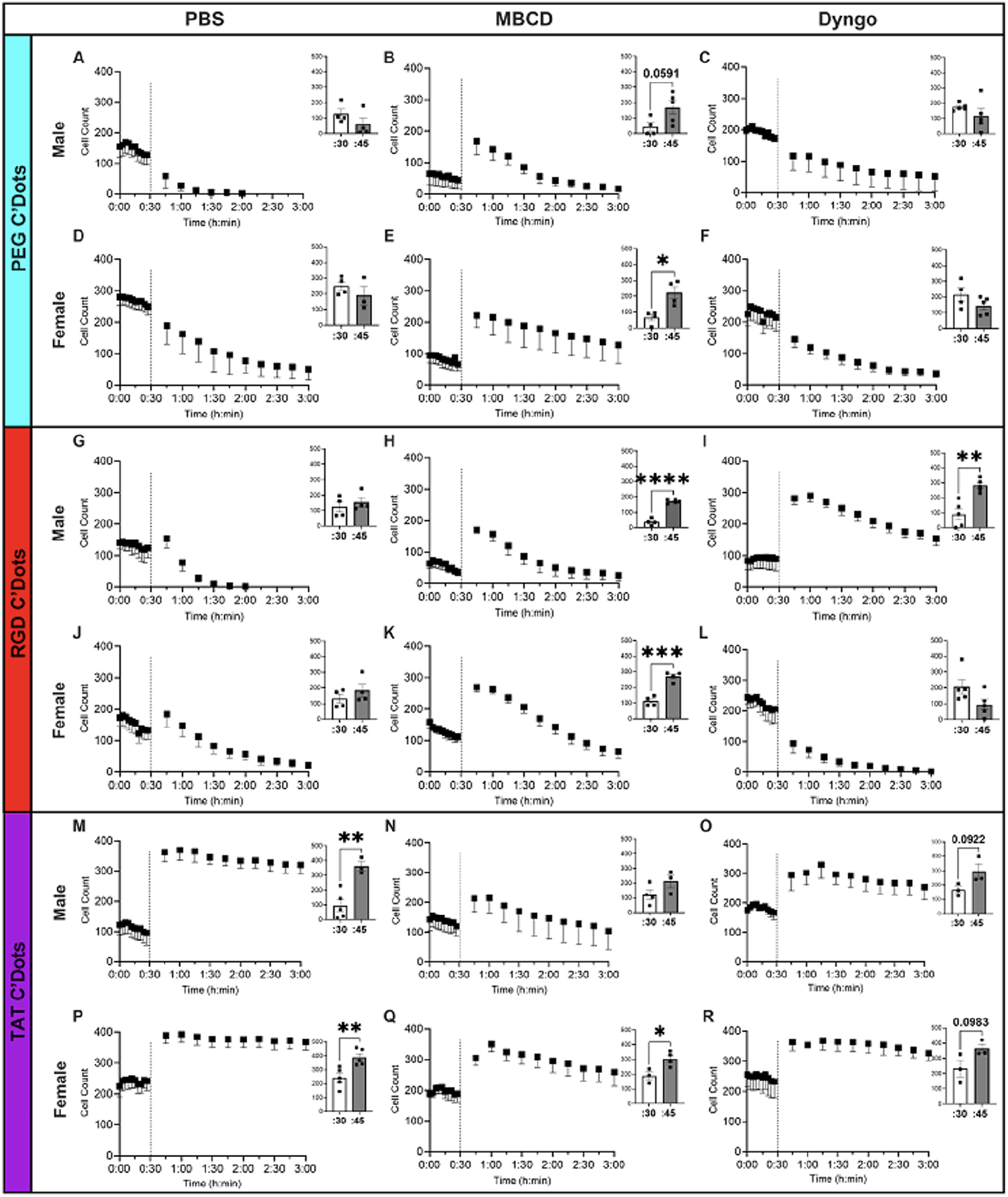
Combination plots of short and long term clearance studies provide insight into how different drugs impact C’Dot kinetics in osteocytes over time. The short study for each drug and C’Dot group is shown in the first 30 minutes of each graph, to the left of the vertical black line. The long study is shown following the vertical black line. PEG C’Dot groups are highlighted in blue on the left (A-F), RGD C’Dot groups are highlighted in red (G-L), and TAT C’Dot groups are highlighted in purple (M-R). For each set of studies, a student’s T-test was performed between the final short timepoint and the initial long timepoint (inset for each graph, *p≤0.05, **p≤0.01, ***p≤0.001, ****p≤0.0001, Error = SEM, n=3-5/group).

**Fig. 18. Figure S10:**
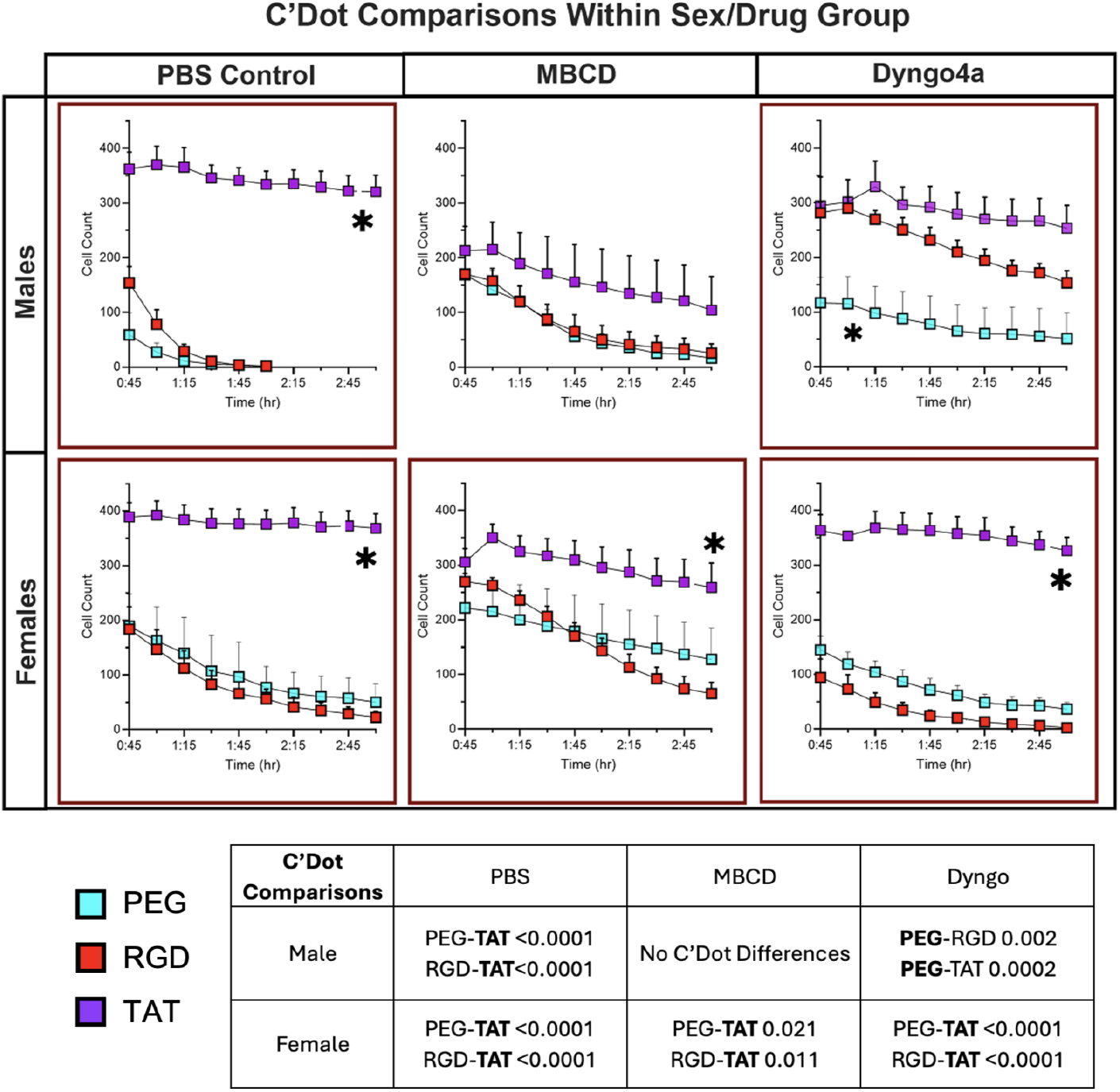
A Mixed Linear Model highlights the impact of C’Dot functionalization on most drug and sex groupings. Plots show mixed model comparisons of C’Dots within drug and sex groups for long incubation clearance data. Comparisons with significant differences are boxed in red. Asterisks denote the group that is distinct from the others within the plot. P≤0.05, Error = ±SEM.

